# Survival Reinforced Transfer Learning for Multicentric Proteomic Subtyping and Biomarker Discovery

**DOI:** 10.1101/2025.01.24.634697

**Authors:** Linhai Xie, Pei Jiang, Cheng Chang

## Abstract

Omics-based molecular subtyping in large-scale and multicentric cohort studies is a prerequisite for proteomics-driven precision medicine (PDPM). However, keeping the subtypes with robust molecular features and significant associations with prognosis among different cohorts is challenging due to the biological heterogeneity and technical inconsistency. Herein, we propose a subtyping algorithm, named Survival Reinforced Patient Stratification (SRPS), to adapt the known subtypes from the discovery cohort to another by simultaneously preserving the distinct prognosis and molecular characteristics of each subtype. SRPS has been benchmarked on simulated and real-world datasets, where it shows a 12% increase in classification accuracy and possesses the best prognostic discriminations. Moreover, based on the calculated subtype significance score, an ‘unpopular’ protein, Peptidylprolyl Isomerase C (PPIC), was identified as the top-1 remarkable protein for subtyping the hepatocellular carcinoma (HCC) patients with the worst prognosis. Eventually, PPIC was experimentally proved to be a pro-cancer protein in HCC, confirming our work as a practice of interpretable machine learning guided biological discovery in PDPM research.

## Introduction

Proteomic subtyping has been broadly applied in oncology research on large cohorts with an important goal of stratifying patients with inconsistent levels of prognosis according to the proteomic features^1–6^. Thorough analysis of differentially expressed proteins between subtypes is able to suggest the potential prognosis biomarkers and drug targets for clinical applications and therefore the research paradigm of proteomics-driven precision medicine (PDPM) was proposed^1^. However, to deliver the molecular stratification of patients from a scientific discovery to a clinical guidance, a fundamental step is to expand its validity to multiple independent clinical cohorts^7–10^ to eliminate the sample bias in a single cohort with a relatively small sample size. Unfortunately, such an expansion of existing molecular subtypes on multiple cohorts is difficult due to unignorable inter-cohort data heterogeneity that stems from both biological divergence and technical inconsistency. Hence, the problem we investigate in this manuscript is how to transfer prognosis-discriminative molecular subtypes from a source cohort to a target cohort which we term as cohort adaptation for molecular subtyping in large-scale and multicentric cohorts.

Commonly applied supervised classification approaches^11,12^ lack the generalization ability between data with biased drifts. Correspondingly, batch effect removal algorithms^13,14^ were proposed for integrating data partially corrupted with batch effects, albeit with the risk of over correction that may lead to unreliable downstream analysis^15^. Another alternative is domain adaptation technique^16–19^ in transfer learning. It can search for data representations that are invariant between cohorts while distinct for each subtype in the source cohort. Nevertheless, its drawback in our scenario is not being able to effectively preserve the prognosis discrimination of subtypes from the source cohort to the target cohort.

As shown in **Figure 1**, we introduce a novel algorithm named Survival Reinforced Patient Stratification (SRPS) for adapting prognostic discriminative patient stratification from the source cohort to a target cohort. SRPS optimizes a classifier to predict the subtype of a patient only based on the molecular profile through two learning paradigms. In the source cohort, it learns the biological features of the known subtypes through supervised learning; while in the target cohort, it learns to maximize the prognostic discrimination between subtypes through reinforcement learning. Hence, SRPS can preserve both the proteomic feature and prognostic significance of each subtype when transferring the subtypes between cohorts.

**Figure 1.**
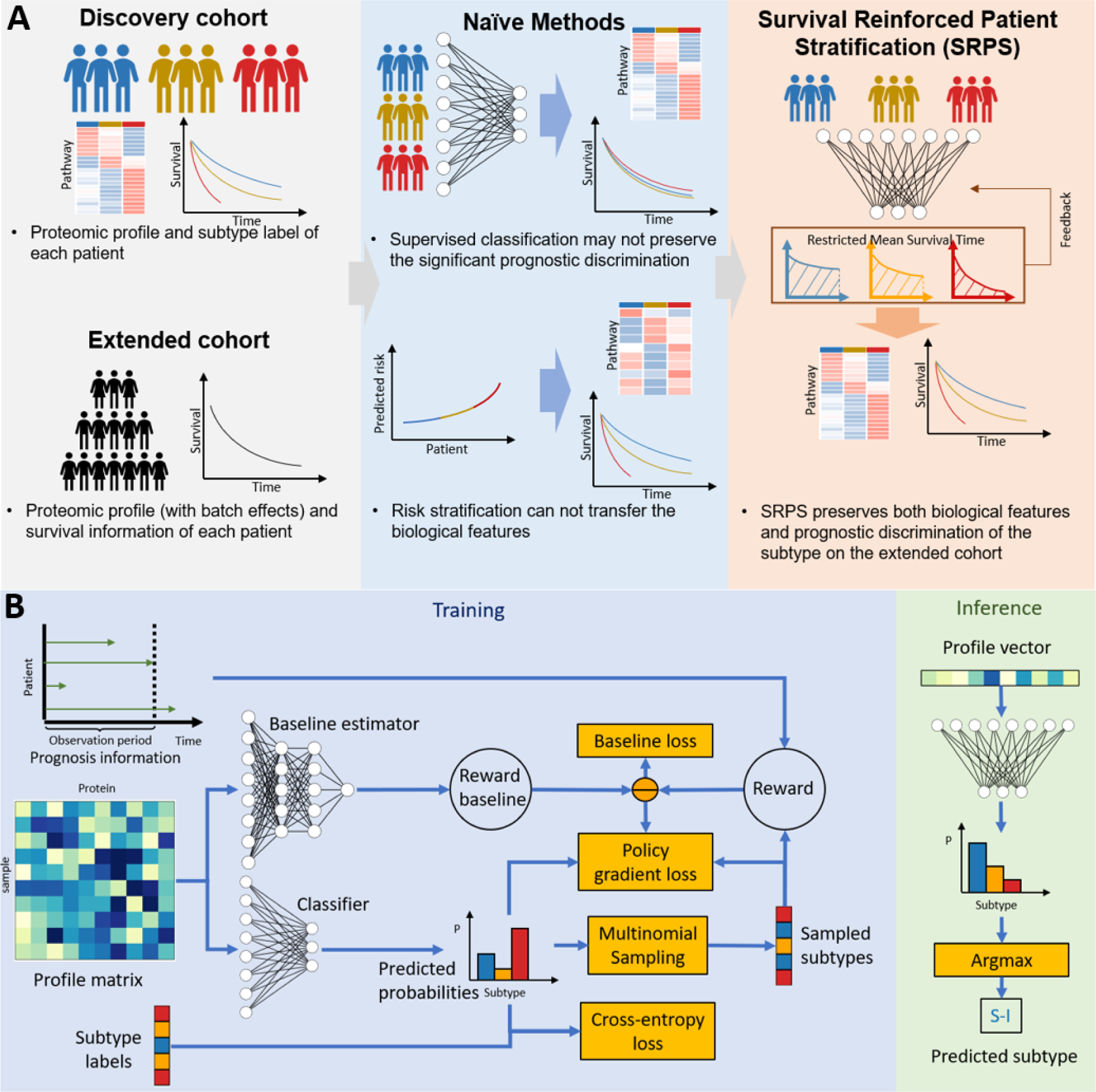
The innovation and the learning strategy of Survival Reinforced Patient Stratification (SRPS) **A**. When transferring subtypes with known biological features and prognostic discrimination from a discovery cohort to an extended cohort with batch effects, naïve methods face significant limitations. Supervised classification often fails to preserve prognostic discrimination, while risk stratification can classify patients but cannot transfer biological features. SRPS addresses these challenges by simultaneously learning from the subtype labels of the discovery cohort and the survival data of the extended cohort. This dual approach enables SRPS to preserve both the biological features and prognostic discrimination of the subtypes. **(b)** During training, SRPS optimizes a classifier to predict patient subtypes based on molecular profiles using two complementary learning paradigms. In the discovery cohort, supervised learning minimizes prediction errors relative to known subtype labels. In the extended cohort, on-policy reinforcement learning maximizes prognostic discrimination between subtypes by optimizing a policy gradient. ⊝ represents element-wise subtraction. In the inference phase, SRPS assigns a predicted subtype to each patient using the trained classifier, as illustrated with the example of a predicted liver cancer subtype (“S-I”).

The performance of SRPS has been benchmarked on both synthetic and realistic datasets (Table S1) against 5 other baseline methods. Experimental results on synthetic datasets corroborate that SRPS outperformed baseline methods by a large margin, 12% higher accuracy and almost 100% greater Log-rank score than the baselines method in average on the target cohort with significant batch effects. As for the real-world datasets, after demonstrating comparable accuracy on the source cohort and ssGSEA similarity between source and target cohorts, SRPS achieves the highest survival discrimination in both overall survival (OS) and recurrence-free survival (RFS) clinical outcomes.

Furthermore, model explanation is always desired for biological and clinical application of machine learning^20,21^ and several strategies^22–24^ have been proposed to interpret complicated models. For this purpose, we designed the network architecture of the classifier with only a single dense layer which enabled a straightforward translation of model parameters with a subtype significance score. By applying the scoring function to models that transferred subtypes between two hepatocellular carcinoma (HCC) cohorts, we observed insightful reasons for the enhanced prognosis discrimination of the classifier. It allocates higher weights on proteins which are originally discriminative in prognosis. More importantly, the scoring function indicated Peptidylprolyl Isomerase C (PPIC) as the most significant protein when subtyping a group of HCC patients with the worst prognosis. It was able to dissect patients into two groups with significantly different prognosis on two HCC cohorts for both OS and RFS, and for the first time, its promotion in proliferation, migration and colony formation of HCC cells were proved on cell lines experiments. These results suggest a potential clinical application of PPIC as a prognostic biomarker.

## Results

### SRPS improves the classification accuracy for prognosis-discriminative subtypes when batch effects exist between cohorts

SRPS aims at improving subtyping accuracy by using the correlation between lifetimes and subtypes through deep reinforcement learning. It lays on two basic assumptions: 1) feature-based subtyping can be improved by exploiting prognosis information; 2) prognosis information can be effectively exploited through deep reinforcement learning.

To prove them, 121 pairs of toy datasets (Figure S1) were simulated, and three different patient stratification strategies are compared in various levels of batch effect between source and target cohorts and the correlation between subtypes and lifetimes (survival correlation). The performance of feature-based stratification strategy based on supervised learning is only affected by the batch effect (**Figure 2A**). On the contrary, the accuracy of survival-based stratification method (Survival), which divides patients with a boundary at 30 of lifetime in the target cohort, is only related to the survival correlation (Figure 2B). SRPS (Figure 2C) shows a more robust performance either with severe batch effects as well as low survival correlation due to the consideration of both features and prognosis information.

**Figure 2.**
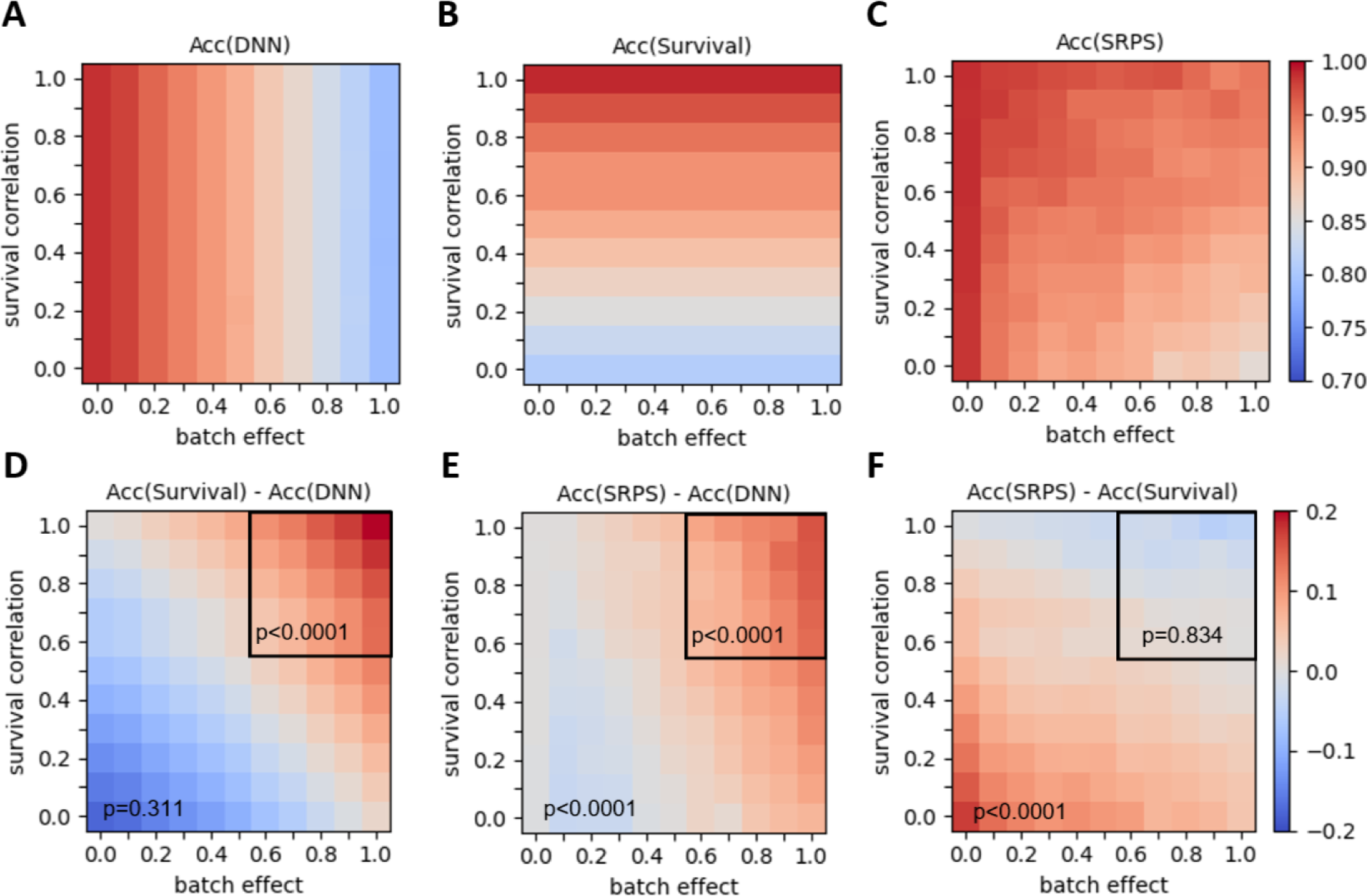
SRPS effectively improves classification accuracy by using prognosis information. Each subfigure displays an accuracy map on 121 simulated toy datasets with various levels of batch effects between source and target cohort and the relative correlation between survival and subtypes (see Methods and Supplementary). In each experiment, 20% samples from target cohort are used for testing. The upper row exhibits the accuracy map of three different stratification strategies on the target cohort. **A**. A feature-based strategy is applied by training a Deep Neural Network (DNN) to predict two subtypes according to features only in the source cohort. **B**. A survival-based strategy directly stratifies patients by thresholding the survival times at 30 months in the target cohort. Note that this strategy is impractical in real since the survival time is unavailable for a certain patient who is still at risk, but here, it is only used as a theoretical reference when such prognosis information could be fully exploited (or accurately predicted) for subtyping. **C**. SRPS uses features for prediction and is adjusted by prognosis information in the target cohort during training phase. **D-F**. The bottom row exhibits the residual accuracy maps between two strategies. The one-tailed Student’s t-test was utilized to exam whether the values in the whole area or the upper-right area are significantly above zeros. Acc means accuracy.

According to the residual accuracy maps between each two strategies, the survival-based method significantly outperforms DNN within the upper right area (*p*<0.0001) with strong survival correlation and large batch effects (Figure 2D), it proves that it is possible to improve classification accuracy in this situation by only considering prognosis information. By contrast, the comparison between SRPS and DNN (Figure 2E) not only shows a significant improvement within upper right area (*p*<0.0001), but also for the whole area (*p*<0.0001). Furthermore, it shows no significant performance inferiority within the upper right area compared to survival-based method (Figure 2F). It proves that SRPS can highly exploit the prognosis information through deep reinforcement learning.

### SRPS outperforms comparative approaches and ablated models in simulated datasets

Here we lend credence to the state-of-the-art (SOTA) performance of SRPS on 2 pairs of simulated datasets (Figure S2) by comparing it with 5 competitive methods, including Random Forests (RF)^11^ and Deep Neural Network (DNN)^12^ for supervised learning, RF combined with Harmony (RFH)^13^ and Deep Adversarial Neural Network (DANN)^16^ for batch effect removal and domain adaptation techniques, semi-deepCLife^25^ for prognosis-guided approach. Then model ablation study was carried out by removing a certain component of reinforcement learning, i.e., SRPS (no baseline), or replacing reinforcement learning with an alternative approach, i.e., SRPS (soft). The evaluation metrics include classification accuracy, ssGSEA similarity, log-rank score and C-index. The mean and standard deviation of all metrics were reported according to a 5-fold cross-validation test which was repeated for 5 times.

#### Comparison with baseline methods

When testing on datasets without batch effect, all these algorithms exhibit excellent prediction accuracy (above 0.9) on both source and target cohorts, showing their generalization ability on samples drawn from the similar distribution as training data (**Figure 3A**). However, their performance degenerates to varying degrees when batch effect exists (Figure 3D).

**Figure 3.**
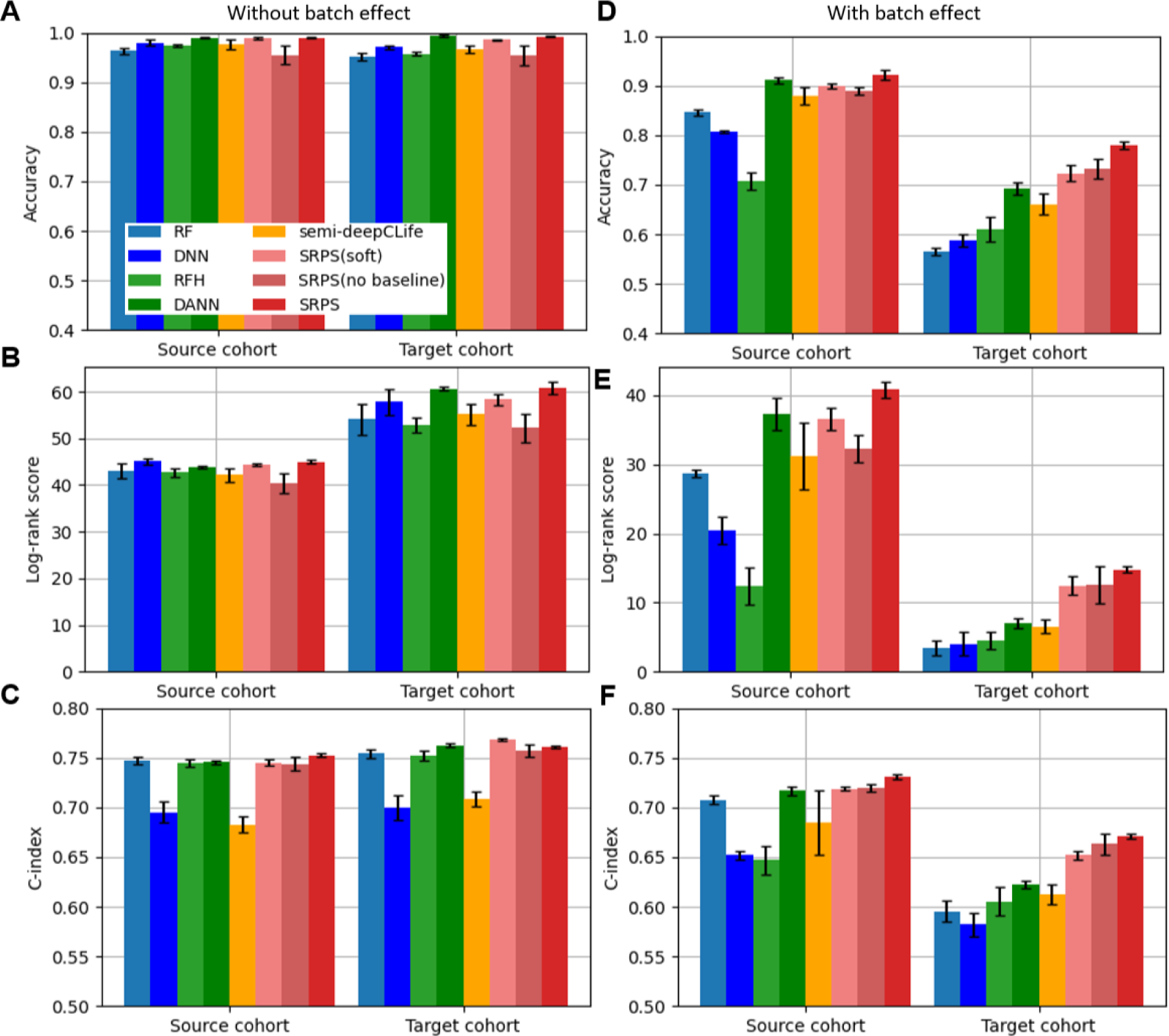
SRPS outperforms comparative approaches and ablated models in simulated datasets. All approaches were firstly benchmarked on two simulated datasets without any batch effect between them **(a-c)**. It was then followed by the benchmarking on datasets with batch effects **(d-f)**. The metrics reported are the mean and standard deviation values of classification accuracy and log-rank score from a 5-fold cross-validation test repeated for 5 times. The competitive methods are random forests (RF), deep neural network (DNN) with supervised learning, RF combined with Harmony (RFH) for batch effect removal, domain adversarial neural network (DANN) for domain adaptation and semi-deepClife for survival guided approach while SRPS (soft) and SRPS (no base line) are ablated models of SRPS.

As the pure supervised learning methods, Random Forests (RF) and Deep Neural Network (DNN) have obvious but reasonable drop of accuracy on the target cohort with a biased distribution drift. RF combined with Harmony (RFH) enhances accuracy compared to the above two approaches on the target cohort, nonetheless, with a non-trivial accuracy drop on source cohort. Its attempts to remove batch effects diminishes the biological feature of each subtype which may partially stem from the fact of not using the labels in the source cohort. Domain Adversarial Neural Network (DANN) is the most powerful baseline approach, though, still falls behind SRPS especially on the target cohort. Although there are other domain adaptation approaches implementing more sophisticated learning mechanisms such as conditional domain adversarial network (CDAN)^26^ and maximum classifier discrepancy (MCD)^17^, they did not achieve higher subtyping accuracy through our observations (Figure S3). Semi-deepCLife, as the most similar competitive algorithm to SRPS, possesses an accuracy only comparative to DANN. There might be two reasons. First, the survival loss of deepCLife does not specify which subtype should have a superior prognosis. This may incur conflicts with the label-based supervision loss during training. The other one could be the lack of random exploration in stochastic gradient descents with a large batch which was required for an accurate estimation of lifetime distributions. Therefore, the straightforward compound of deepCLife and supervised learning is not an effective solution. In contrast, SRPS outperforms all competing methods by a large margin when batch effects exist between cohorts. More specifically, it improves the target domain accuracy by 12% compared to the second-ranked model, DANN.

The diagnostic discrimination ability of each model is also illustrated. For the case without batch effect (Figure 3B and 3C), all methods have high log-rank score and C-index, since they have shown accurate prediction in subtypes which are strongly correlated to prognosis, both for source and target cohorts. On the contrary, their diagnostic discrimination decreases remarkably on the target cohort with large batch effects (Figure 3E and 3F). As expected, SRPS enhances the log-rank score by more than 100% compared to the best baseline model.

#### Model ablation study

Here we verified the contribution of two critical reinforcement learning-related components. We firstly eliminated the reward baseline estimator which is important for gradient stabilization in many policy gradient-based reinforcement learning algorithms^27^. The gradients estimated by REINFORCE^28^ are calculated from random samples and always cause noisy rewards. As shown in Figure. 3, the accuracy, log-rank score and C-index on the target cohort reported for SRPS (no baseline) have higher standard deviation than the full SRPS method, indicating the high variance during the learning procedure of the ablated model.

The other ablation model is to replace the entire reinforcement learning component with soft relaxation as in deepCLife which is an alternative for dealing with the non-differentiable component in a neural network. The drawback of applying soft relaxation in our situation is possibly due to the dilemma of batch size selection during training as mentioned above. As a result, this design corrupted the performance of SRPS (Figure 3).

### SRPS outperforms comparatives approaches in real-world cohort adaptations of proteomic subtypes

In this section, we evaluated all approaches in two different cohort adaptation scenarios with three hepatocellular carcinoma (HCC) cohorts and a lung adenocarcinoma (LUAD) cohort to demonstrate the SOTA performance of SRPS. Each cohort is provided with proteomic profiles of all samples and the time and status of clinical outcomes including overall survival (OS) and recurrence-free survival (RFS).

Jiang et al.’s cohort^1^ recruited 101 early stages (0-A) Chinese HCC patients according to Barcelona Clinic Liver Cancer (BCLC) staging system^29^. According to the original paper of Jiang et al. cohort, all patients in this cohort were classified into 3 subtypes with different proteomic features and prognosis as visualized in Figure S4, which will be transferred to other target cohorts as the known subtype. Gao et al.’s cohort^2^ involved 159 Chinese patients with broader BCLC stages, additionally including stage B and stage C. Xing et al.’s cohort^30^ included 152 Chinese patients with the same range of BCLC stages as Gao et al.’s cohort. Xu et al.’s cohort^5^ recruited 103 Chinese patients with another type of cancer, i.e., LUAD. According to Figure S5, the proteomic profile matrices of these cohorts exhibit large inter-cohort heterogeneity.

#### Transfer HCC subtypes to other HCC cohorts with broader cancer stages

The first experiment is to transfer the proteomic subtyping of HCC patients from Jiang et al.’s cohort to Gao et al.’s cohort and Xing et al.’s cohort respectively. **Figure 4A** and B show the testing results. Except RFH, all approaches share similar prediction accuracy on the source cohort. Deep learning-based methods (DNN, DANN, semi-deepCLife and SRPS) have comparatively higher ssGSEA similarities of the classified subtypes between source and target cohorts, indicating that neural networks learn more generalizable features of each subtype. A similar trend remains for the Log-rank score on OS. Pure supervised learning or domain adaptation techniques such as DNN and DANN do not perform well in this metric mainly due to the reason that the RFS distributions of all subtypes in the source cohort are originally not significantly different. Therefore, this attribution is naturally inherited by its prediction on the target cohort. Semi-deepCLife was also designed to improve discrimination through exploiting two kinds of survival information together. It performed comparatively to SRPS in C-index. In the end, SRPS exhibits powerful discrimination ability in terms of both OS and RFS with comparable classification accuracy on source cohort and ssGSEA similarity between two cohorts. Exemplary results of SRPS among 5 repeated tests on Gao et al.’s cohort and Xing et al.’s cohort are visualized in Figure S6 and S7. It is worth noting that the clinically acceptable prognosis discrimination should reach a log-rank score above −*log*10(0.05)≈1.3.

**Figure 4.**
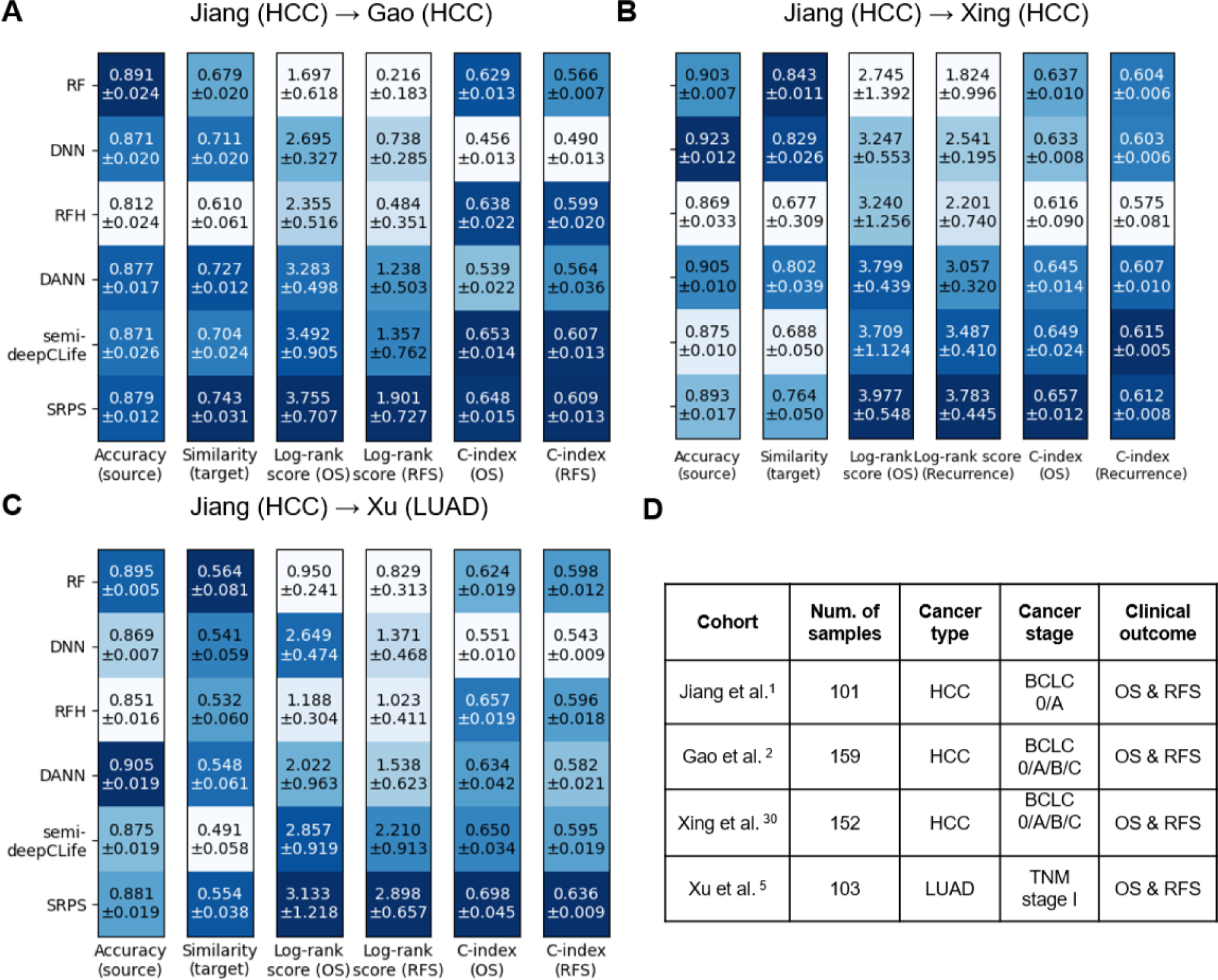
SRPS outperforms comparatives approaches on real-world cohort data. **A**. Subtypes discovered from Jiang et al.’s cohort were transferred to Gao et al.’s cohort. **B**. Subtypes discovered from Jiang et al.’s cohort were transferred to Xing et al.’s cohort. **C**. Subtypes from Jiang et al.’s cohort were transferred to Xu et al.’s LUAD cohort. Metrics include the accuracy in the source cohort, ssGSEA similarity of subtypes between the source and target cohorts, log-rank score, and C-index for overall survival (OS) and recurrence-free survival (RFS). Larger values represent better performance, as indicated by deeper colors in each column. **D**. Summary of the four real-world cohorts, including cancer type, number of samples, cancer stage, and clinical outcomes evaluated.

#### Transfer HCC subtypes to a LUAD cohort

Next, we attempted to transfer subtypes from Jiang et al.’s HCC cohort to Xu et al.’s LUAD cohort (Figure 4C). Among all methods, SRPS again exhibits the highest prognostic discrimination in OS and RFS, demonstrating its superiority in optimizing prognosis discrimination of subtypes. However, we find that the ssGSEA similarities of all methods are lower than the similarities when transferring between two HCC cohorts. It is reasonable since the onset of cancer at different organs can have drastically heterogeneous biological processes. Thus, HCC patients and LUAD patients with consistent prognosis are unlikely to share similar proteomic patterns between their tumor samples. This observation infers that although SRPS enhances the probability of detecting biologically and prognostically consistent subtypes between two cohorts, it compromises by reducing the biological consistency, when the original subtypes are unlikely to exist in the new cohort naturally. Its discoveries obey the underlying biological facts. An exemplary result of SRPS among 5 repeated tests on Xu et al.’s cohort can be visualized in Figure S8.

### PPIC is identified as a significant protein through model interpretation with subtype significance score

The interpretability of models is of importance in biomedical research, where a common solution is to keep the model as simple as possible. Therefore, the classifier utilized in SRPS for real-word proteomic experiments is a single layer neural network with a parameter matrix shaped as *S* × *D* and is activated by a soft-max layer for probabilistic outputs. Based on this simple model, we proposed a subtype significance metric named Δweight to indicate the contribution of each protein to classify patients into a specific subtype. It is worth noting that the scale of the expression of each protein has been processed with z-score normalization so that the contribution of each protein for subtyping does not depend on the range of its original expressions. The total number of Δweight values equal to the number of values in the model parameter matrix. A detailed definition of Δweight is provided in Methods.

Based on the experimental results to transfer HCC subtypes from Jiang et al.’s cohort to Gao et al.’s cohort, we investigated the weight matrix of all 25 classifier models of SRPS (5 random repeats × 5 folds) and focused on the subtype S-III of HCC patients since it indicates the worst prognosis after surgery. The subtype significance score averaged among all 25 models was used in the following analysis to alleviate the influence of random initialization. Additionally, since the subtype significance score Δweight is defined based on the model weights which vary during the training procedure, we carry out stability experiments. Figure S9 informs that the rank of the top-10 proteins based on the average subtype significance score becomes stable quickly within the training phase. Besides, the relationship between its ranking stability and the number of models required has been discussed in Figure S10.

#### SRPS classifies patients with prognosis-discriminative features

We first evaluated the correlation between the subtype significance score of SRPS classifier model and prognosis discrimination of each protein in Gao et al.’s cohort. The prognosis discrimination of each protein was measured by the coefficient of a univariate Cox regression model and the correlation was estimated by the linear regression. Consequently, in terms of OS, the Δweight of SRPS model has a correlation (r=0.60) similar to that of DNN (r=0.55) (**Figure 5A**) mainly because that DNN has already achieved good discrimination in OS (Figure 4A), albeit, SRPS shows much higher correlation (r=0.55 vs r=0.32) in terms of RFS (Figure 5A), as it enhances the discrimination of RFS by a large margin (Figure 4A). This result informs us that SRPS improves the prognostic discrimination of subtypes by allocating higher weights on features that are more discriminative in prognosis during training than the method without exploiting survival information.

**Figure 5.**
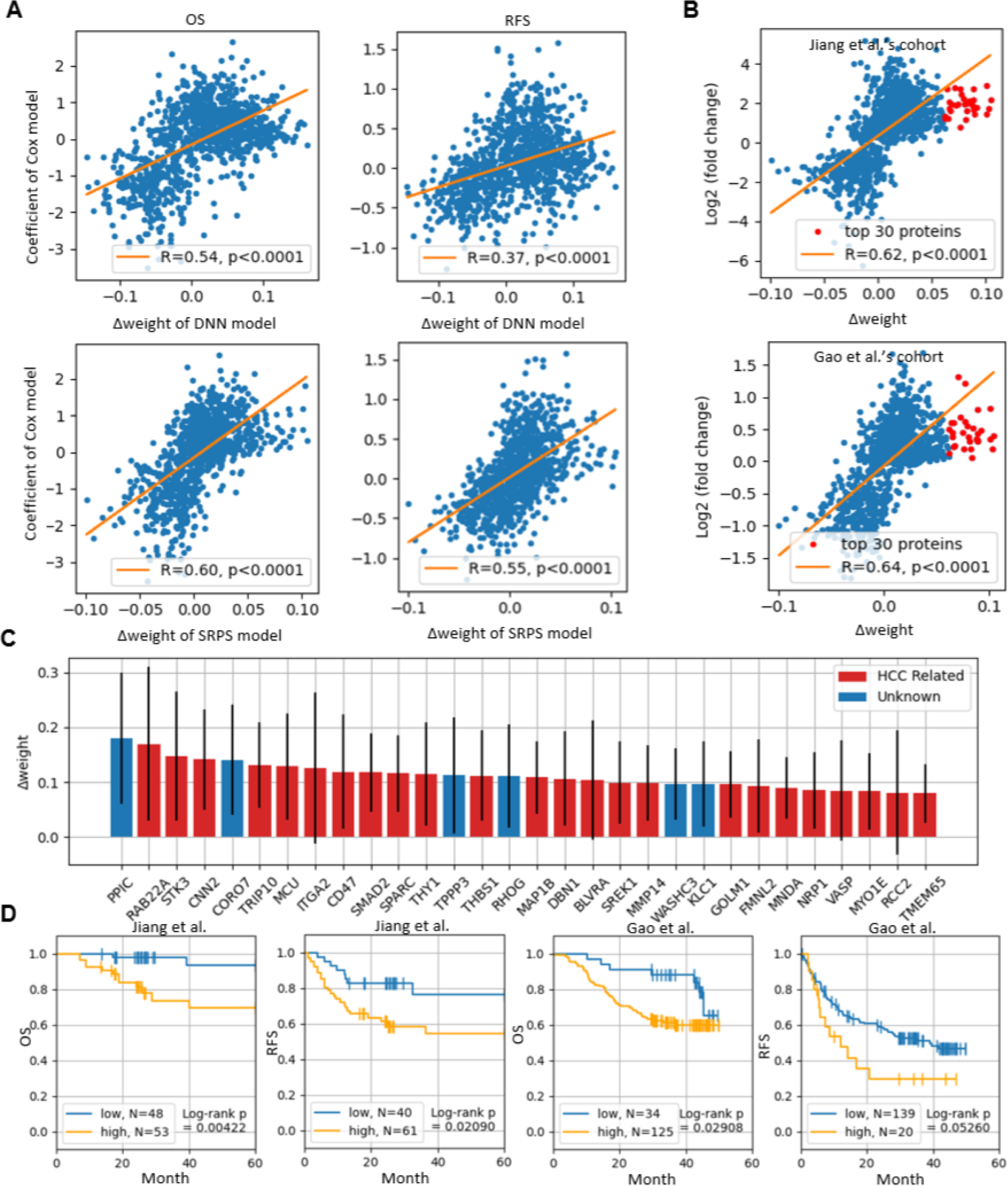
PPIC was identified as a significant protein through model interpretation with subtype significance score. **A**. The correlation between the prognostic discrimination (the coefficient of a univariate COX model) and the subtype significance score (Δweight) of each protein in the model (DNN and SRPS) in terms of OS and RFS. **B**. The correlation between the subtype significance score of each protein in the model and the corresponding log2(fold change) value on Jiang et al.’s cohort and Gao et al.’s cohort. Red points indicate proteins with top-ranked significance scores. **C**. The mean and standard deviation of the significance score of the top-30 proteins and whether they have been formally reported to be HCC-related in published literatures. **D**. The prognostic discrimination power of PPIC in terms of OS and RFS on Jiang et al.’s cohort and Gao et al.’s cohort. The threshold for dividing high and low expression groups in each subfigure is independently set to be the value with the highest log-rank test significance.

#### SRPS encourages the discovery of proteins different from differential expression analysis

A similar and common approach to identifying signature proteins for each subtype is Differential Expression Analysis (DEA) which usually serves as a clue for biomarker discovery^1,2^. Hence, we calculated correlation between Δ*weight* and log2 fold change value of protein expression for S-III in each cohort where the latter is an essential metric for DEA (Figure 5B). It exhibits a positive correlation and, more importantly, obvious discrepancy in proteins with peak values, i.e., top 30 proteins with the highest subtype significance scores do not possess correspondingly pinnacle log2 fold change values. This observation infers a potential ability of subtype significance scores to uncover valuable proteins which might be ignored by classical DEA-based approaches.

#### PPIC identified by SRPS is an under-explored HCC-related protein and is discriminative in prognosis

We further investigate the 30 top-ranked proteins about whether they or their corresponding genes have been formally reported with an association with HCC in research publications (Figure 5C). As a result, 24 of 30 proteins were found with literature evidence to support their relevance with HCC progress, interpreting that SRPS classifies patients by exploiting biological meaningful features. Among the six under-explored HCC-related proteins, limited functional evidence has been reported for **CORO7** (Coronin-7) and **WASHC3** (WASH complex subunit 3) in cancer types. However, **PPIC** (Peptidyl-prolyl cis-trans isomerase), **TPPP3** (Tubulin polymerization-promoting protein family member 3), **RHOG** (Rho-related GTP-binding protein RhoG), and **KLC1** (Kinesin light chain 1) have been shown to play roles in cancer progression, such as cell proliferation and metastasis, in other cancer types (Table S2). These findings suggest their potential involvement in HCC. More importantly, we found that PPIC protein, which corresponds to the highest subtype significance score has received little attention in HCC-related research, thus triggering our deeper investigation into that protein.

The survival analysis in Jiang et al.’s cohort and Gao et al.’s cohort for both OS and RFS by stratifying patients according to the expression of PPIC shows its prognosis discrimination ability in these two clinical outcomes (Figure 5D). The threshold was independently grid-searched in each case and the value resulting in the largest significance of log-rank test was selected. As shown in results, PPIC is discriminative in prognosis and can potentially be developed into a biomarker of HCC patients with poor survivals.

### PPIC functions as a pro-cancer protein in HCC

Although PPIC was reported to be highly expressed in several cancers, its role in HCC remains elusive. To determine the potential functions of PPIC in HCC, we knocked down PPIC expression using two different siRNAs targeting PPIC in HCC cell lines. As shown in **Figure 6**A, the PPIC mRNA expression was significantly reduced in cells treated with the siRNAs targeting PPIC respectively. Notably, as shown in Figure 6B-C, PPIC knockdown profoundly inhibited cell proliferation and colony formation in two HCC cell lines-Huh7 and MHCC-LM6, which suggests that PPIC promotes HCC progression. Similarly, knockdown of PPIC also significantly inhibited cell colony formation in other two HCC cell lines-Hep3B and MHCC-97H in Figure S11A-B. Besides, Figure S11C shows that cell migration was strongly inhibited by PPIC knockdown in Huh7 cells. Thus, these results indicate that PPIC functions as a pro-cancer protein in HCC, which is consistent with the results from SRPS, and the molecular mechanism behind it needs further investigation.

**Figure 6.**
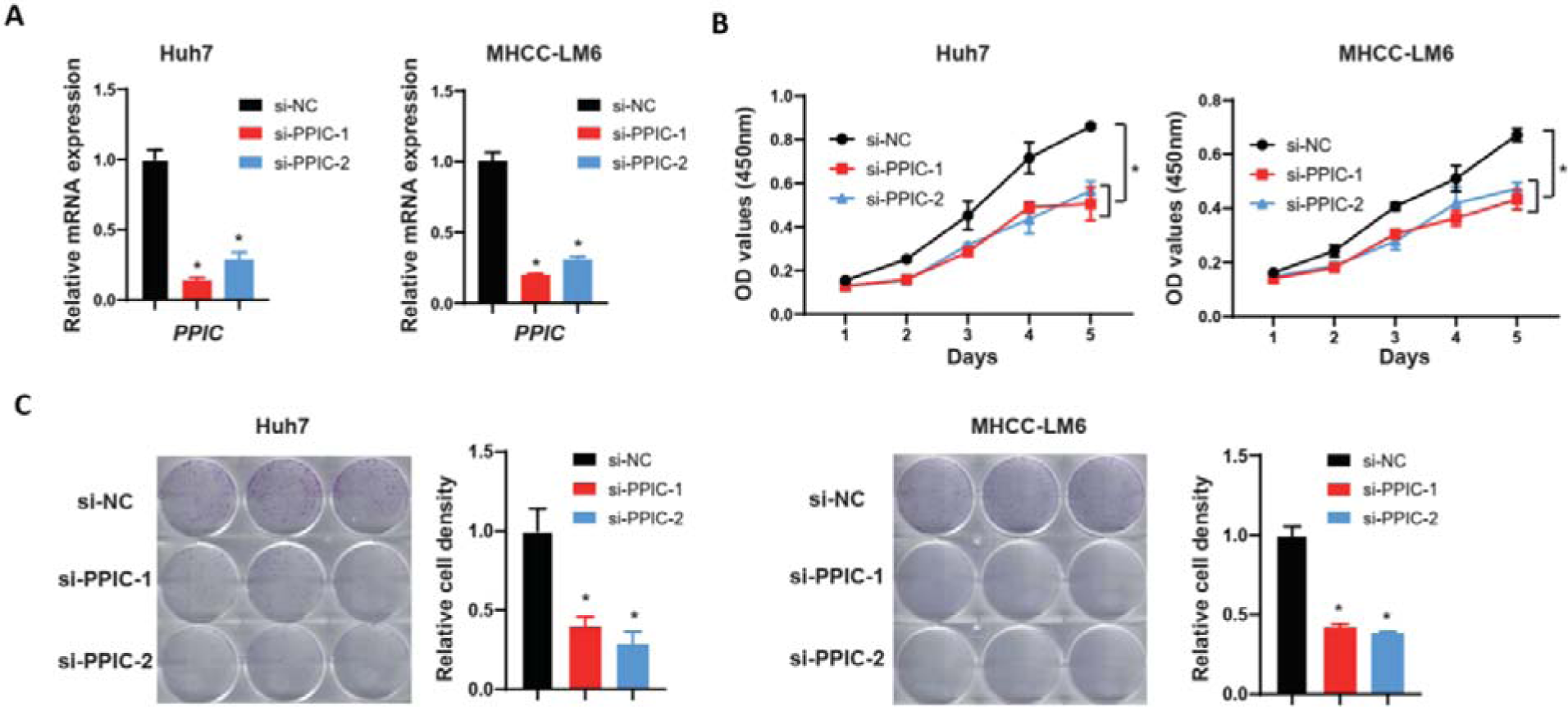
PPIC promotes cell proliferation and colony formation in HCC cell lines. **A**. Relative mRNA expressions of *PPIC* after the indicated treatment. **B**. Cell proliferation curves determined by cell viability assay after the indicated treatment. **C**. Resulting cell colonies (left) and the related statistical graphs (right) after the indicated treatment. Data were presented as mean±SEM. *, ***p***<0.05* (Two-tailed Student’s t-test).

## Discussion

Multicenter study is essential in clinical oncology research to eliminate the biological and environmental biases of the sampled population. Therefore, to deliver the practical value of PDPM to clinical applications, expanding and validating the discoveries from a single cohort to multiple ones becomes an inevitable request. However, since proteins are the minimal functioning units of the human body system, proteome is a more real-time observation of the disease progress compared to genome and transcriptome, therefore it exhibits large data heterogeneity between cohorts.

We proposed the SRPS algorithm to adapt known proteomic subtypes from a labeled source cohort to the unlabeled target cohort. Unlike existing batch effect removal or domain adaptation techniques which seek for indistinguishable representations between batches/domains with the risk of ignoring essential biological difference, SRPS unifies the classification features between cohorts by preserving the prognosis discrimination of subtypes which is important for clinical applications. Through experiments on simulated and real-world data, we demonstrated the effectiveness of guiding the subtyping with survival information through reinforcement learning.

Deep reinforcement learning has shown great success in optimizing a neural network with a self-defined reward function by encouraging desired behaviors and penalizing undesired ones without explicit labels as in gaming^31^, robotics^32^ and drug design^33^. It also provides a solution to estimate gradients of non-differentiable^34^ evaluations of the predictions, alongside soft approximation^25,34^/relaxation^35^ as an alternative. By comparing SRPS with semi-deepCLife or SRPS (soft), we have witnessed higher performance in prognosis discrimination of the former option and, in general, this strategy can be applied to any objective due to the high degree-of-freedom in designing the reward function. For instance, it is possible to reinforce the ssGSEA similarity of the subtypes between two cohorts by incorporating it into the reward function.

Since survival information has been proved effective in guiding the learning process of subtyping, sophisticated architectures of the neural network become unnecessary to achieve acceptable subtyping performance. As a result, it enables the use of a single dense layer in the classifier which brings simplicity in model interpretation and alleviates the problem of overfitting. Consequently, PPIC was discovered as a critical protein in identifying HCC patients with poor prognosis.

Here, we demonstrate that PPIC is required for HCC cancer cell proliferation and colony formation, indicating its pro-cancer functions. Though PPIC was reported to be involved in endoplasmic reticulum redox homeostasis and hepatic stellate cell activation^36,37^, its key molecular functions and direct substrates in HCC remain to be further elucidated. Thus, our results showed that PPIC is a powerful prognostic marker for HCC and the molecular revelation of its pro-cancer mechanism might contribute to HCC treatment.

Despite its promising results, SRPS has certain limitations. Currently, SRPS is restricted to transferring known subtypes from a source cohort to target cohorts and does not have the capability to discover unified new subtypes across multiple cohorts. In future work, we aim to address this limitation by integrating SRPS with unsupervised clustering methods, enabling the discovery of novel subtypes from diverse datasets.

Additionally, while we validated the role of PPIC in this study, other subtype-significant proteins identified by SRPS have yet to be experimentally examined. Future efforts will focus on validating these proteins through functional experiments, which may further elucidate their biological relevance and clinical utility in cancer subtyping.

## Methods

### Survival Reinforced Patient Stratification

As illustrated in Figure 1B, the training framework of SRPS consists of a subtype classifier and a reward baseline estimator. The classifier predicts the subtyping probabilities of patients given the proteomic profile matrix. For data with subtype labels in the source cohort, the predicted probability as well as the provided ground truth is used to calculate a cross-entropy loss. By minimizing this objective, the classifier can learn the principle proteomic pattern of each subtype and stratify patients according to it.

More specifically, given x_i_ ∈ R^D^ to represent the D dimensional proteomic profile vector of sample *i*∈*I*, where *I* is the patient set, a prediction vector is calculated as:

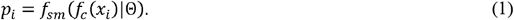

Each element 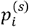 in *p*_*i*_ represents the probability of the patient *i* to be identified as the subtype *s* ∈ {1, 2,…, S}, supposing *S* different subtypes in total. *f*_*sm*_(·) and *f*_*c*_ (·)|Θ) stand for the soft-max activation function and the neural network classifier parameterized with Θ respectively. With the one-hot encoded label *y*_*i*_ in source cohort, the cross-entropy loss is calculated as:

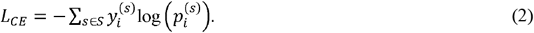

Meanwhile, the classifier adjusts its prediction with the guidance of prognostic information of each patient on the target cohort through an on-policy reinforcement learning algorithm, REINFORCE^28^. It samples a subtype prediction ŷ _*i*_ of patient *i* from a multinomial distribution parameterized by the predicted subtype probability *p*_*i*_ as:

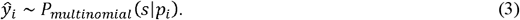

Such a random sampling strategy encourages the exploration of better classification boundaries and is only used for training. The subtype prediction for inference is deterministic with an argmax operation given predicted probabilities *p*. Then, for a specific subtype *s* we calculate the restricted mean survival time (RMST) *τ*_*s*_. This calculation is based on two aspects related to each patient within the population of subtype *s*: one is the lifetime *t*_*i*_ of the patient, and the other is the observation of the patient’s survival status, which we denote as *d*_*i*_.

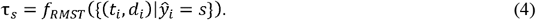

The detailed calculation of *f*_*RMST*_ is provided in latter sections. All RMST values formulate a reward *r* to assess the prognosis discrimination of the predicted subtypes as:

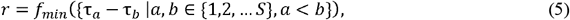

where *f*_*min*_ outputs the minimal value from a set and without the loss of generality, we use a smaller number to annotate the subtype with a better prognosis which is known a priori from the source cohort. However, since such behavior evaluation with rewards is usually noisy, a reward baseline is required for stabilization^27^ which includes an input-dependent component *b*_*i*_ and an input-independent component *b*_*c*_ as:

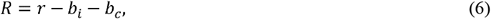

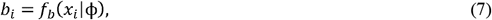

where *f*_*b*_ represents a neural network predictor with ϕ as parameters and both types of baseline components are adjusted by minimizing the baseline loss:

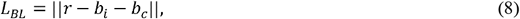

where ∥·∥ is the L2 norm. Afterwards, the predicted subtype probabilities, the sampled subtypes and the baseline-stabilized rewards produce an approximation of the policy gradient^27,28^ as:

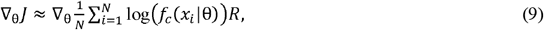

where *N* is the number of patients. It regulates the classifier with the aim of improving its prognostic discrimination. The above objective can be implemented with a cross-entropy loss given the predicted probability *p* and the sampled prediction *ŷ* as:

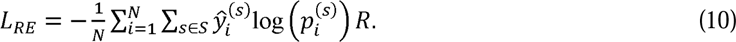

The final loss function for training the classifier is:

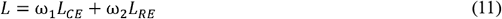

where ω_l_ and ω_2_ are coefficients between 0 and 1. Note that above procedure describes the training phase of the classifier and there is no need for survival information during inference phase.

The calculation of RMST is simple, but it requires the predicted subtyping results for all the patients in the mini-batch, which is a discrete variable and is undifferentiable. Hence, we exploit the on-policy reinforcement learning algorithm REINFORCE by taking ^34^ as a reference to approximate the true gradients and keep an end-to-end training procedure. Furthermore, the reason for using an on-policy algorithm is that it is simpler for implementation compared to an off-policy algorithm which needs more elaborate tuning on the replay buffer and the exploration strategy.

### Simulated dataset generation

121 pairs of toy datasets were created to examine two basic assumptions about SRPS. The original pair of datasets (top in Figure S1) involves a source and a target cohort with identical features *x*_*origin*_ whose values ranges between [0,1] and the feature dimension is 20. Normally distributed random noise *σ*_*noise*_ was added to the features as batch effects between the source and target cohorts. The mean of the noise vector was uniformly sampled from the interval of [0,1] independently along each dimension and the standard deviation was set to 0.2. The resulted feature vector is *x*_*origin*_ + ασ_*noise*_ where we gradually increase the noise weight α from 0 to 4 with a step of 0.4, corresponding to the lowest batch effect scale 0 to the strongest scale 1 respectively as shown in Figure 2. The survival time of all patients was strictly divided according to the subtype labels where S-I patients all survive longer than 30 months and S-II patients die before 30 months without any censored value for simplicity. To decrease the relative survival correlation between the subtype and survival times which leads to a weaker prognostic discrimination of subtypes we randomly swap the survival time of patients between two subtypes. The swapping ratio ranges from 0 to 0.2 with a step of 0.02, corresponding to the strongest relative correlation 1 and weakest relative correlation 0 respectively as shown in Figure 2.

The simulated datasets for benchmarking (Figure S2) are generated with the R package Splatter^38^, with the function splatSimulate. We generate two sets of datasets with/without batch effects. In each dataset we simulate 1000 total samples with 3 different subtypes which are divided into 2 batches. The batch effect between batches are controlled with two parameters batch.facLoc and batch.facScale which are both set to 0.01 for no batch effects and 2 for large batch effects. The feature size of all samples is set to 1000. The survival data are generated with the R package Simsurv^39^, following the Weibull distribution with the coefficients as (λ = 0.005, γ = 0.5). We carried out two experiments to validate the reliability of the simulated data. Figure S12 demonstrates that the distributional divergence between the simulated data and a real-world HCC cohort is as close as the divergence between two real-world HCC cohorts. Besides, Figure S13 indicates that the difficulty of predicting patient prognosis, measured with C-index by using a random survival forest model, between the simulated data and a real-world cohort is also comparable.

### Accuracy maps on the toy datasets

We generated 121 pairs of toy datasets to examine the influence of batch effects between the source and target datasets, as well as the correlation between subtype labels and survival times in the target cohort (termed survival correlation). Details of the dataset generation process are described in the **Simulated dataset generation** section. The 121 pairs of datasets are a Cartesian product of 11 batch effect levels and 11 survival correlation levels. Each cube in the accuracy maps in Figure 2 represents the result of experiments conducted on a specific pair of source and target datasets, with batch effect and survival correlation levels indicated by the axes of the sub-figures. The x-axis represents increasing batch effect levels, and the y-axis represents increasing survival correlation levels.

For each experiment, the evaluation metric is prediction accuracy, which we define as:

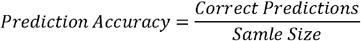

where accuracy is calculated on 20% of the samples in the target cohort not used for model training. This definition is consistently applied to all experiments involving synthesized datasets.

Three accuracy maps (Figure 2A-C) were constructed for different subtyping strategies:

1. **Figure 2A (Feature-based strategy):** A Deep Neural Network (DNN) trained on features in the source cohort predicts two subtypes.
2. **Figure 2B (Survival-based strategy):** Patients in the target cohort are stratified by thresholding survival times at 30 months. This strategy, while theoretical due to its reliance on unavailable survival data for at-risk patients, serves as a reference for fully exploiting prognosis information.
3. **Figure 2C (SRPS):** Our proposed algorithm.

To further compare these strategies, three residual accuracy maps (Figure 2D-F) were generated via element-wise subtraction between accuracy maps. For example, Figure 2D was created by subtracting the accuracy of the survival-based strategy from that of SRPS across all 121 dataset pairs.

### Real-world proteomic data preprocessing

The raw proteomic expression matrix of each cohort is preprocessed before experiments. Let the rows and columns of the matrix indicate samples and proteins respectively. The preprocessing contains following steps: **1)** Each row of the profile matrix was normalized with the quantile normalization; **2)** All the proteins expressed in less than 30% of the samples were filtered; **3)** All missing values were imputed with zero; **4)** Each column was normalized with Z-score normalization; **5)** We use the intersection of the proteins simultaneously identified in both source and target cohorts; **6)** The resulted protein sets in two scenarios are then intersected with the signature protein sets suggested by the paper of source cohorts respectively.

Consequently, when transferring the subtypes from Jiang et al.’s cohort to other two HCC cohorts or to Xu et al.’s LUAD cohort we used the signature proteins suggested from the literature^1^ and the feature dimension of proteomic profile matrices is 1097. (Table S1)

### Restricted mean survival time (RMST)

Given a series of paired survival times *T*= *T*_l_, *T*_2_,…, *T*_n_ and status (dead or not) *D*= {*D*_l_, *D*_2_,…, *D*_n_} of *n* patients, we can firstly estimate a Kaplan-Meier (KM) curve which consists of successive survival probabilities *s*_*t*_at each time within the time interval *t*= {0, *t*_l_, *t*_2_,…, *t*_*max*_}. *t*= *t*_0_, we have *S*_0_ = 1, and the consecutive probabilities can be extrapolated as follows:

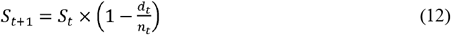

where *d*_*t*_ counts the number of subjects died between [*t, t* + 1] and *n*_*t*_ counts the number of subjects living at the start. Note that subjects who are lost are regarded as censored and are not involved in *n*_*t*_. The RMST is calculated as the area under this KM curve within a restricted time interval, i.e. from 0 to 60 months in our experiments.

Using the restricted mean survival time (RMST) is because of its simplicity and directionality. When compared with other divergence metrics such as the Kullback-Leibler, the Maximum-Mean Discrepancy (MMD) and the Kuiper test that only tell the distance between the survival distribution of two subtypes as mentioned in ^25^, the difference between the RMST of two subtypes is simple to calculate and clearly indicates which one holds a better prognosis. Therefore, it can largely boost the learning efficiency from the prognosis information.

### Training settings of SRPS

The classifier of SRPS consists of only one linear layer with l1 regularization (coefficient=ϕ) and dropout where drop rate is set to ρ_*su*_ for supervised learning and ρ_*rl*_ for reinforcement learning. The output dimension equals the number of subtypes. The baseline estimator consists of 3 layers with *n*_*h*_ neurons in each hidden layer with sigmoid function as activation except the output layer. The output dimension is 1.

Both models are optimized with Adam optimizers given learning rate α_*rl*_ and α_*bl*_ respectively. The models are trained with the full batch for 10000 episodes with/without using early stop (*Es* = *true*/*false*) which save the model with the highest Δ*RMsT* on validation set.

Hyper-parameters are manually selected according to the performance on validation set within the possible combinations of following settings: *n*_*h*_ = {10,100}, ω_l_ = {0.1,0.01,0.001}, ω_2_ = 1, ϕ= 10, 10, ρ_*su*_ = {0,0.3,0.8}, ρ_*rl*_ = {0,0.2}, *Es* = {*true, false*}. Given all possible combinations of hyper-parameters, we select the best model with different criteria on simulated and real-world datasets respectively. For testing the model with simulated data, we apply the validation accuracy as the selection criteria. When testing the model on real-word datasets, we pick the model with the highest Log-rank score (OS) and a ssGSEA similarity score no less than 0.5 on the validation set, since there is no subtyping label for the target dataset.

The model and training process was implemented with Python and Tensorflow library and was deployed on a server equipped with an InteI XII Gold 6242R CPU and a Quadro GV100 GPU. Each run of training lasted about 2 minutes.

### Single sample gene enrichment score analysis (ssGSEA) similarity

For each patient in the real-word proteomic datasets, functional enrichment analysis^40^. was performed using the ssGSEA algorithm implemented in the R package GSVA^41^. The quantile normalized protein expression matrix was imported to GSVA to calculate the ssGSEA scores for each gene set with at least ten overlapping genes. The interested gene set was provided in the literature^15^. After obtaining the enrichment scores of interested gene set of each patient, we calculated the average enrichment score for each subtype and estimated the similarity between cohorts with the cosine similarity. The final similarity score between the two cohorts is the average similarity of all subtypes.

### Log-rank score and C-index

The log-rank score was defined as – *log*_10_(*P*) based on the P value of a log-rank test^42^, which indicates how likely all groups having the same lifetime distribution. A higher score means better prognosis discrimination. Concordance index (C-index)^43^ calculates the fraction of pairs of subjects for which the model predicts the correct order of survival while also considering censoring. 0.5 indicates random predictions and 1.0 means perfect predictions. It was calculated by checking the concordance between the survival time and probability difference between subtypes with the best and worst survivals, i.e., *p*_*i*_^(l)^ − *p*_*i*_^(3)^ where *p*_*i*_ is the predicted probability vector of patient *i* and its first and third dimensions represent the probability of being S-I (best prognosis) and S-III (worst prognosis) respectively.

### 5-fold cross-validation

All benchmarking results (except for testing on toy datasets) are reported based on a 5-fold cross-validation experiment repeated for 5 times with different random seeds. More concretely, for each repeat, we set the random seed and evenly split the data into 5 folds. Then a loop starts by selecting 3 of the folds for training, one for validation and the rest for testing. It finishes until all 5 folds have been used as the test fold respectively and the testing results of all 5 folds are regarded as the final testing result of the current repeat.

### Implementation of competitive algorithms

RF algorithm is an ensemble of tree predictors where all trees vote for the most likely class prediction. We used the implementation from the python package sklearn^44^ with the function RandomForestClassifier where we set the max depth to 10 and trained it as a classification model under the supervision of subtyping labels provided in the source cohort.

DNN is a neural network that consists of a hidden layer, a dropout layer and a prediction layer and was trained purely in a supervised manner. It was implemented with Tensorflow library in python. The hidden dimension and dropout rate were set to 20 and 0.8 respectively to avoid overfitting. The cross-entropy loss and Adam optimizer were utilized. The learning rate was set to 0.01 and the model was trained for 2000 epochs.

Harmony is an algorithm designed to remove batch effects on single-cell data by projecting data from various batches into a batch-invariant representation space through soft clustering^13^. Then, samples can be classified with RF within this space without being affected by batch effects. We firstly did embeddings of the raw profile matrix with PCA to obtain the low-dimensional representations (100 or 10 dimensions) of the data and then applied the python implementation of Harmony algorithm from package harmony-py with its default parameter settings.

DANN is a counterpart solution in machine learning and computer vision area which contents with the domain adaptation task. There are some more SOTA algorithms other than DANN for image domain adaptation^17,26,45^, howbeit, we found they could not lead to any improvement potentially due to the lack of effective neural network structure in encoding omics data (Figure S2). The DANN was implemented with python tensorflow library. It includes an encoder *z*= *t*_*en code*_(*x*), a task classifier *ŷ* = *f*_*task*_ (*z*) and a domain classifier 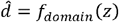. On the one hand the task classifier was trained to minimize the cross-entropy loss between the subtype predictions and labels, on the other hand, the domain classified was optimized to correctly predict the domain of the sample, i.e. the sample was drawn from the source or target cohorts. The critical trick was to insert a layer that can reverse the gradients between the domain classifier and the encoder which encourages the encoder to learn domain-invariant features to confuse the domain classifier. The number of layers in the encoder and classifiers were tuned with two options (1 and 2). The hidden dimension of the encoder and classifiers was set to 20 and 10 respectively. The learning rate was set to 0.01, and the dropout rate was tuned with two options (0 and 0.3).

DeepCLife^25^ is the deep learning-based approach that solves an unsupervised clustering problem with survival information. For fairness, its classifier is optimized by minimizing the original survival loss on target cohort together with the supervised loss on source cohort, which we term as semi-deepCLife. We consider semi-deepCLife to be the most relevant baseline method to SRPS. We reimplemented the tensorflow version according to the original opensource code available at (https://github.com/PurdueMINDS/DeepLifetimeClustering) which is based on the pytorch library. The parameters tuned for semi-deepCLife are learning rate= {0.01,0.001}, l1 regularization= {10^−5^, 10^−4^}, dropout rate= {0.3, 0.8} and weight of survival_loss= {0.01, 0.05}.

### Ablated models of SRPS

The first ablated model is to remove the baseline predictions of rewards in SRPS. It changes the Eq. 6 to *R*= *r* and stops optimizing the baseline predictor. The second ablation is to replace the reinforcement learning approach into a soft relaxation strategy as used in deepCLife. The main difference is that instead of applying the Kuiper test p-value based loss function, we here calculate and maximize a soft version of Δ*RMsT* for optimization. More concretely, when calculating *d*_*t*_ and *n*_*t*_ in Eq. 12 for a specific subtype group, the count of each subject is weighted by the probability of the subject belonging to this subtype. Such an operation keeps the entire tensor graph differentiable and can be directly optimized through back-propagation algorithm.

### Definition of subtype significance score Δ*weight*

Given the parameter matrix of the classifier in SRPS as Θ∈ *R*^*S×D*^, the logits vector *l*∈ *R*^*S*^ based on an input profile vector *x*∈ *R*^*D*^is formulated as *l*= Θ*x* and the predicted probability vector is:

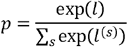

where *D* and *S* are the profile dimension and the number of subtypes respectively and *l*(^s^)indicates the *s*^th^ element in the vector *l*. Unlike the random sampling based on predicted probabilities during training, the subtype prediction in the inference stage is deterministic with argmax operation as shown in Fig.1. In other words, for *a, b* ∈ {1, 2, …, *S*} where *b* ≠*a*, the predicted subtype *s* will be *a* iff *p*(^*a*^) <*p*(^*a*^)⇒*l*(^*a*^)<*l*(^*b*^)⇒Θ(^*a*^)*x* < Θ(^*b*^)*x* holds true for any possible *b*, where Θ(^*a*^)represents the *a*^th^ row vector of Θ. Therefore, when we only focus on a specific feature *x* (^*i*^),*i* ∈ {1,2,…, *D*}, we can approximately have:

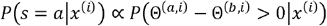

where *b* ∈ {1, 2,…, *S*} and *b* ≠*a*. Θ(^*a, i*^)represents the element at *a*^th^ row and *i*^th^ column of Θ. The above equation informs us that the probability of predicting the given profile as subtype *a* when only considering a single protein expression *x*(^*i*^)is proportional to its corresponding weight Θ(^*a, i*^)and inversely proportional to the weight of other subtypes. Accordingly, we give the following definition about Δ*weight*:

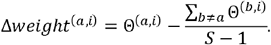

Since this metric is defined based on the model weights and can possibly be affected by the random initialization of the model, we utilized the averaged value of multiple models to indicate significant proteins and the convergence of this approach during the training process was shown in Figure S9.

### Cell culture and siRNA transfection assay

HCC cell lines-Huh7, Hep3B, MHCC-LM6 and MHCC-97H were cultured in Dulbecco’s Modified Eagle Medium (DMEM) supplemented with 10% fetal bovine serum (FBS), 100 U/ml penicillin, and 100 mg/ml streptomycin, and the cells were incubated at 37C with 5% CO_2_. Cells were seeded in 6-well plates before transfection. When cell density approached 50%, cells were transfected with the indicated siRNAs twice at 24-hour interval using TurboFect Transfection Reagent (Thermo). Cells were collected for mRNA expression assays 48h after transfection. The siRNA sequences targeting *PPIC* are as follows:

si-PPIC#1: GCUCUAGCAACAGGAGAGA;

si-PPIC#2: CUCGAUCAUCAACAGUGGC.

### mRNA expression assay

Total RNA was extracted using TRIzol Reagent (Invitrogen) according to the manufacturer’s instructions, and the cDNA was obtained using HiScript® III All-in-one RT SuperMix Perfect for qPCR (Vazyme). The gene expression was measured by real-time PCR (qPCR) using Taq Pro Universal SYBR qPCR Master Mix (Vazyme) with a Bio-Rad CFX96 instrument. Statistical graphs were generated using GraphPad Prism 8.3.0 software. The qPCR primers used in this paper are as follows:

*ACTIN*-Forward: 5’-CACCATTGGCAATGAGCGGTTC-3’;

*ACTIN*-Reverse: 5’-AGGTCTTTGCGGATGTCCACGT-3’;

*PPIC*-Forward: 5’-CTGCTGCTACCTCTCGTGC-3’;

*PPIC*-Reverse: 5’-GCCAATCACAATTCTGCCAACA-3’.

### Cell viability and colony formation assays

For cell viability assay, after the indicated treatments, approximately 5×10^3^ cells per well were seeded into 96-well plates. In the next five days, the cells were treated with CCK8 kit (Aqlabtech) and the OD values (450 nm) were obtained using Spark® Microplate Reader (TECAN) per day. For cell colony formation assay, after the indicated treatments, approximately 5×10^3^ cells per well were seeded into 6-well plates. After 12 days, cells were fixed with methanol and then incubated with Crystal Violet Staining Solution (Beyotime) for 1 h. Finally, the cells were washed with PBS three times and the resulting colony pictures were taken by a camera. Statistical graphs were generated using GraphPad Prism 8.3.0 software.

### Cell migration assay

After the indicated treatment, 1×10^5^ cells were resuspended in 200 μL DMEM without FBS and seeded onto the upper chamber of a transwell filter with 8-µm pores. The lower chamber contained 500 μL DMEM with 20% FBS. 48h later, cells on the underside of the filter were fixed with methanol and then incubated with Crystal Violet Staining Solution (Beyotime) for 1h. After washing, photos were taken by microscope equipped with a digital camera. Statistical graphs were generated using GraphPad Prism 8.3.0 software.

## Data availability

The code to generate synthetic datasets and the processed datasets of real-world HCC and LUAD cohorts are available at https://github.com/PHOENIXcenter/SRPS. The raw proteomic profiles and clinical information can also be downloaded from following pages: https://www.nature.com/articles/s41586-019-0987-8 for Jiang et al.’s HCC cohort, https://www.cell.com/cell/fulltext/S0092-8674(19)31003-7 for Gao et al.’s HCC cohort, https://www.cell.com/cell-reports-medicine/fulltext/S2666-3791(23)00509-8 for Xing et al.’s HCC cohort and https://www.cell.com/cell/fulltext/S0092-8674(20)30676-0 for Xu et al.’s LUAD cohort.

## Code availability

The full code in R and Python is available via GitHub at https://github.com/PHOENIXcenter/SRPS.

## CRediT author statement

**Linhai Xie:** Conceptualization, Data curation, Formal analysis, Methodology, Software, Validation, Visualization, Writing – original draft. **Pei Jiang**: Investigation, Methodology, Resources, Visualization, Writing – original draft. **Cheng Chang**: Conceptualization, Formal analysis, Funding acquisition, Supervision, Writing – review & editing.

## Supplementary material

Supplementary material is available at *Genomics, Proteomics & Bioinformatics* online (xxxx).

## Acknowledgements

This work was supported by the National Key Research and Development Program of China [2021YFA1301603 and 2020YFE0202200], the National Natural Science Foundation of China [32088101 and 82203601], and CAMS Innovation Fund for Medical Sciences (CIFMS) [2019-I2M-5-063].

