## Supplementary Information for "Survival Reinforced Transfer Learning for Multicentric Proteomic Subtyping and Biomarker Discovery" (from another version of this preprint: version 1, 2025-01-27)

**Table S1 Datasets used in experiments**

| Dataset | Data type | Num. of feature used | Clinical outcome | Num. of samples |
| --- | --- | --- | --- | --- |
| 121×2 toy datasets | Simulated data | 20 | Survival Time | 250 / dataset |
| 2 splatter datasets (without batch effect) | Simulated data | 1000 | Survival Time | 500 / dataset |
| 2 splatter datasets (With batch effect) | Simulated data | 1000 | Survival Time | 500 / dataset |
| Jiang et al.’s HCC cohort | Real-world proteomics data | 1097 | OS & RFS | 101 |
| Gao et al.’s HCC cohort | Real-world proteomics data | 1097 | OS & RFS | 159 |
| Xing et al.’s HCC cohort | Real-world proteomics data | 1097 | OS & RFS | 152 |
| Xu et al.’s LUAD cohort | Real-world proteomics data | 1097 | OS & RFS | 103 |

HCC: hepatocellular carcinoma; LUAD: lung adenocarcinoma; OS: overall survival; RFS: recurrence-free survival.

**Table S2 Results of functional experiments** **in other cancer types from literature regarding the six under-explored HCC-related proteins discovered by SRPS**

| **Protein name** | **Abbrev.** | **Cancer type** | **Results of functional experiments** | **Ref.** |
| --- | --- | --- | --- | --- |
| Peptidyl-prolyl cis-trans isomerase C | PPIC | Melanoma | PPIC was identified as a promoter of melanoma progression, enhancing cell invasiveness while concurrently suppressing CD8+T cell activation. | ^1^ |
|  |  | Gastric cancer | PPIC as a gene highly expressed in CAFs was closely associated with NCT (Neoadjuvant chemotherapy) resistance in gastric cancer. | ^2^ |
| Coronin-7 | CORO7 | \ | \ | \ |
| Tubulin polymerization-promoting protein family member 3 | TPPP3 | Glioblastoma | TPPP3 promote epithelial-mesenchymal transition via Snail1 in glioblastoma. | ^3^ |
|  |  | Endometrial carcinoma | TPPP3 knockdown displayed evident suppression in cell proliferation, migration and invasion in vitro and in vivo. | ^4^ |
|  |  | Breast cancer | Silence of TPPP3 suppresses cell proliferation, invasion and migration via inactivating NF-κB/COX2 signal pathway. | ^5^ |
|  |  | Non-small-cell lung cancer | Knockdown of TPPP3 suppresses proliferation and induces apoptosis in non-small-cell lung cancer. | ^6^ |
|  |  | Non-small-cell lung cancer | TPPP3 promotes cell proliferation, invasion and tumor metastasis via STAT3/ Twist1 pathway in non-small-cell lung carcinoma. | ^7^ |
|  |  | Lewis lung carcinoma | Stable knockdown of TPPP3 by RNA interference in Lewis lung carcinoma cell inhibits tumor growth and metastasis. | ^8^ |
|  |  | Colorectal cancer | Knockdown of TPPP3 inhibits cell proliferation and invasion. | ^9^ |
|  |  | Nasopharyngeal carcinoma | TPPP3 overexpression inhibits cell proliferation and invasion in nasopharyngeal carcinoma. | ^10^ |
| Rho-related GTP-binding protein RhoG | RHOG | Salivary adenoid cystic carcinoma | RhoG overexpression promoted SACC-83 cell migration and invasion through activating Rac1. | ^11^ |
|  |  | Glioblastoma | The miR‑124‑3p interacted with RhoG, inhibited RhoG expression and suppressed glioblastoma cell proliferation and migration. | ^12^ |
|  |  | Glioblastoma | siRNA-mediated depletion of RhoG inhibits colony formation and invasion of glioblastoma cells. | ^13^ |
| WASH complex subunit 3 | WASHC3 | \ | \ | \ |
| Kinesin light chain 1 | KLC1 | Breast cancer | Loss of KLC1 inhibited cell metastasis and stem cell markers expression. | ^14^ |
|  |  | Glioma | KLC1-ROS1 fusion upregulated cell proliferation, invasion, and chemoresistance when compared to wild-type ROS1. | ^15^ |

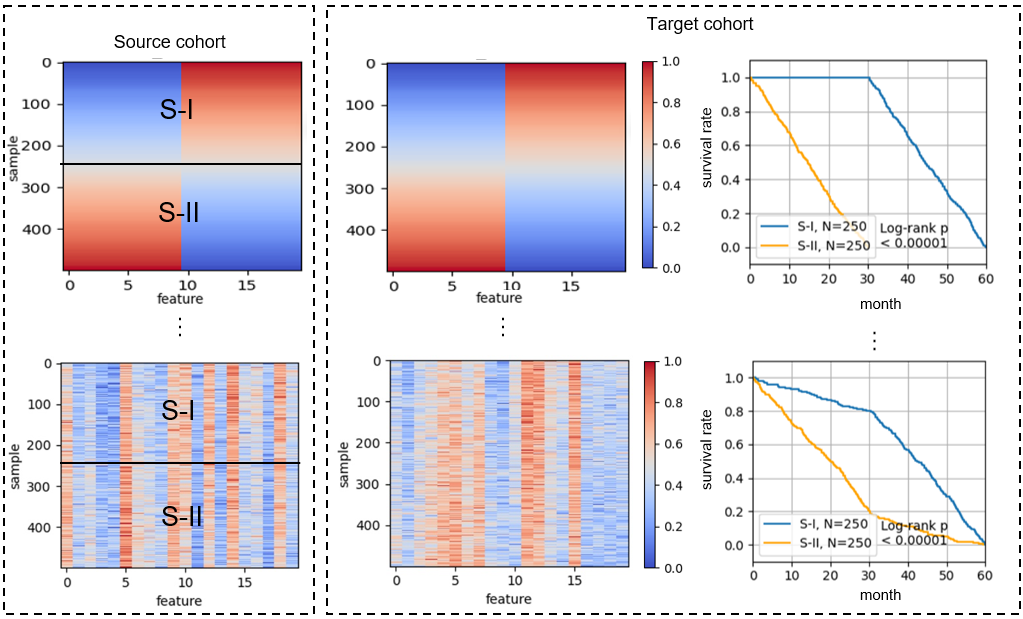

**Figure S1 The visualization of 121 toy datasets used for proof of concepts**

The feature in source and target cohorts are originally identical without batch effects and the survival time between two subtypes are strictly separated. The survival time of S-I samples is all longer than 30 months while the survival time of S-II samples are all shorter than 30 months. We polluted the feature with 10 different levels of random noise to simulate batch effects. Meanwhile, we also randomly swapped the survival time between two subtypes with 10 different swapping rates to decrease the correlation between subtypes and survival time. It finally resulted in 11×11=121 datasets.

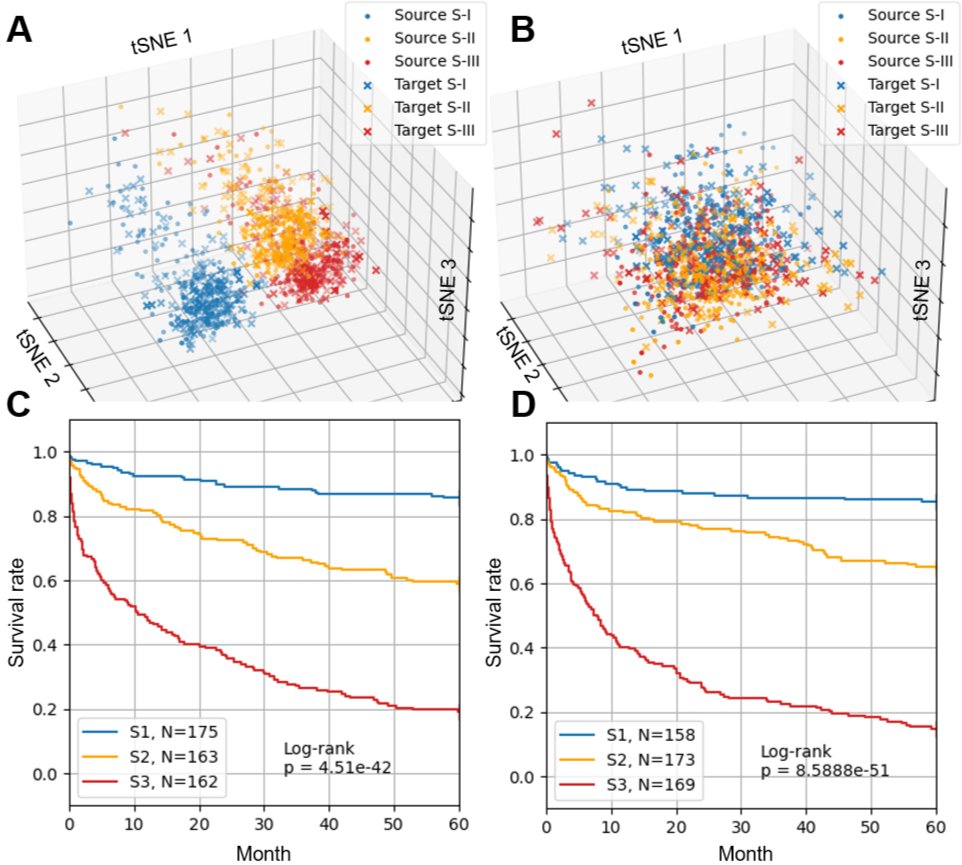

**Figure S2 The visualization of simulated datasets for benchmarking**

The 3D t-SNE visualization of simulated data with different levels of batch effects (**A**-**B**) and the Kaplan-Meier curves of the synthetic prognosis of each subtype in the target cohort (**C**-**D**). Each color indicates a subtype while dots and crosses represent samples from source cohort and target cohort respectively.

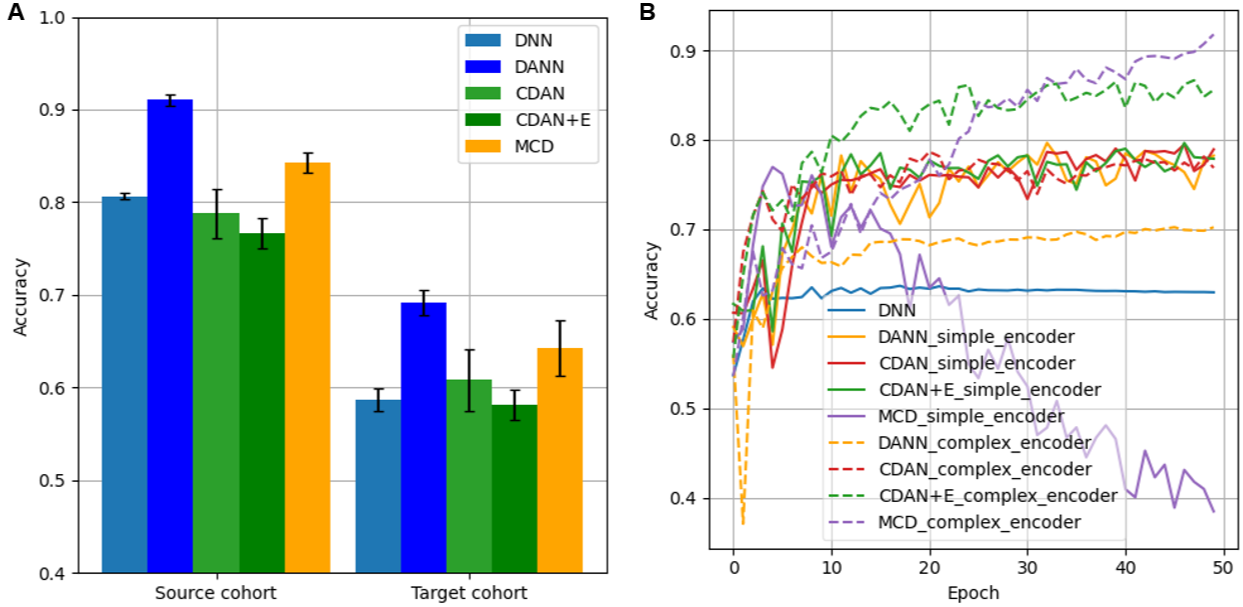

**Figure S3 More advanced domain adaptation techniques did not improve proteomic subtyping**

**A.** The subtyping accuracy (mean±standard deviation) of the supervised learning method (DNN) and domain adaptation approaches, including domain adversarial neural network (DANN), conditional domain adversarial network (CDAN), the improved version of CDAN with entropy (CDAN+E) and maximum classifier discrepancy (MCD), on simulated datasets with batch effects. It is shown that more advanced approaches such as CDAN(+E) and MCD performed worse than DANN. **B.** The accuracy curves of different domain adaptation approaches on MNIST (greyscale) and MNIST-M (colorful) datasets^16^. Domain adaptation approaches were tested with a simple encoder (two layers of CNN, solid lines) or a complex one (three layers of CNN together with batch normalization modules, dashed lines). It proved that the performance gain of CDAN(+E) or MCD largely stems from the exploitation of a powerful image encoder. Since only fully connected neural layers were applied as the encoder for processing omics data in our experiments, DANN was selected for benchmarking with domain adaptation techniques.

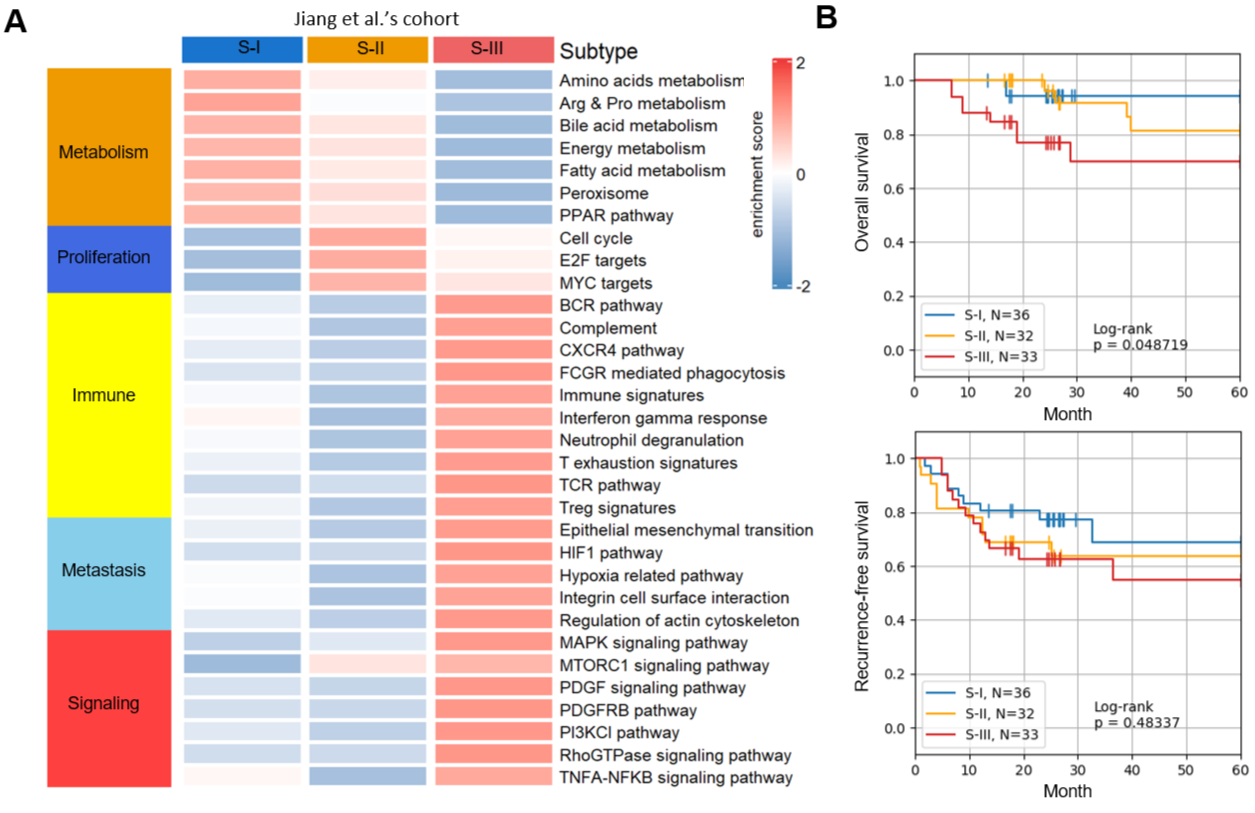

**Figure S4 The visualization of Jiang et al.’s cohort**

**A.** The pathway enrichment scores of the originally discovered subtypes. **B.** The Kaplan-Meier curves of overall survival and recurrence-free survival of the originally discovered subtypes.

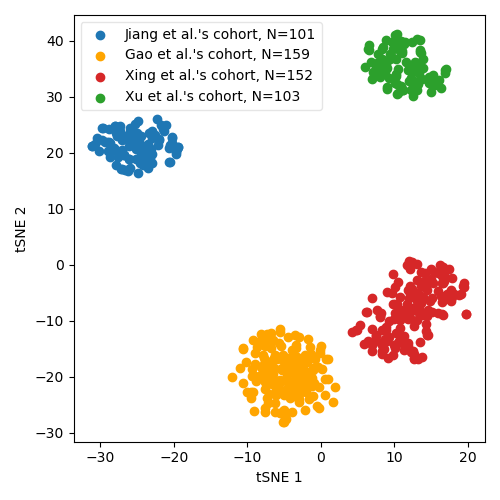

**Figure S5 The data heterogeneity between four real-world proteomic datasets**

The proteomic profile matrix of four real-world cohorts is visualized with 2D t-SNE. Samples belonging to each cohort aggregated into separatable clusters, demonstrating unignorable inter-cohort heterogeneity.

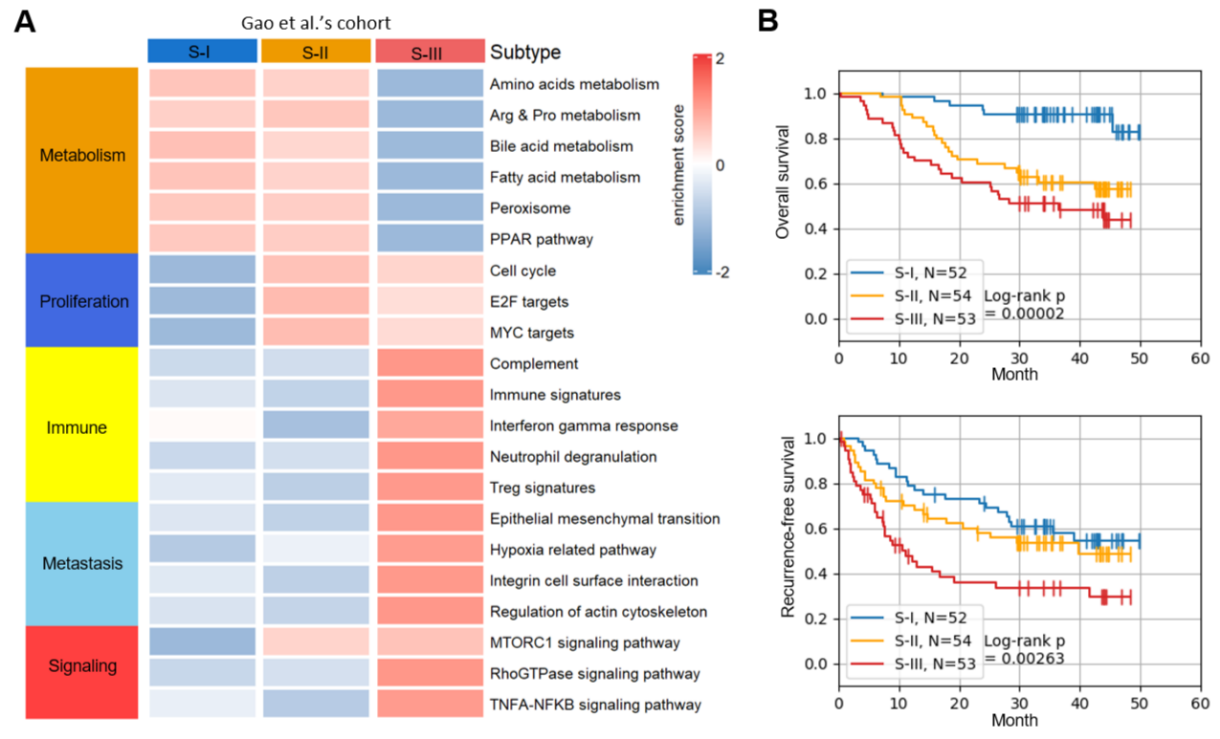

**Figure S6 The visualization of subtyping results on Gao et al.’s cohort**

**A.** The pathway enrichment scores of the SRPS transferred subtypes. **B.** The Kaplan-Meier curves of overall survival and recurrence-free survival of the SRPS transferred subtypes.

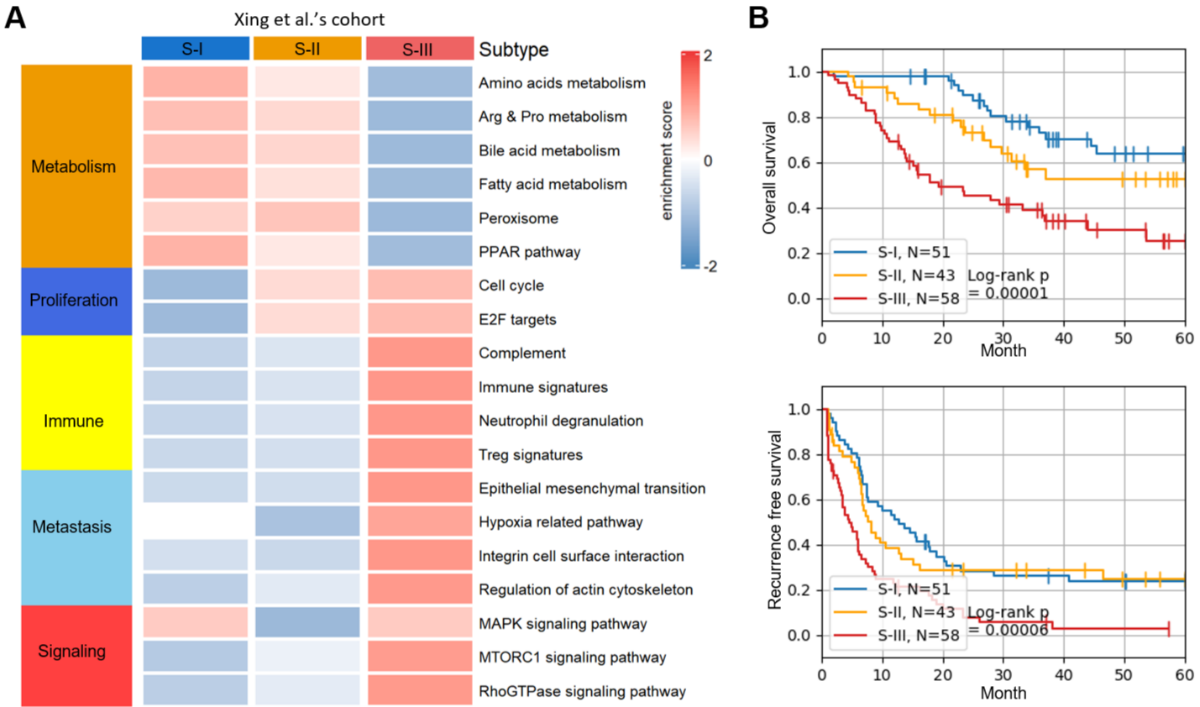

**Figure S7 The visualization of subtyping results on Xing et al.’s cohort**

**A.** The pathway enrichment scores of the SRPS transferred subtypes. **B.** The Kaplan-Meier curves of overall survival and recurrence-free survival of the SRPS transferred subtypes.

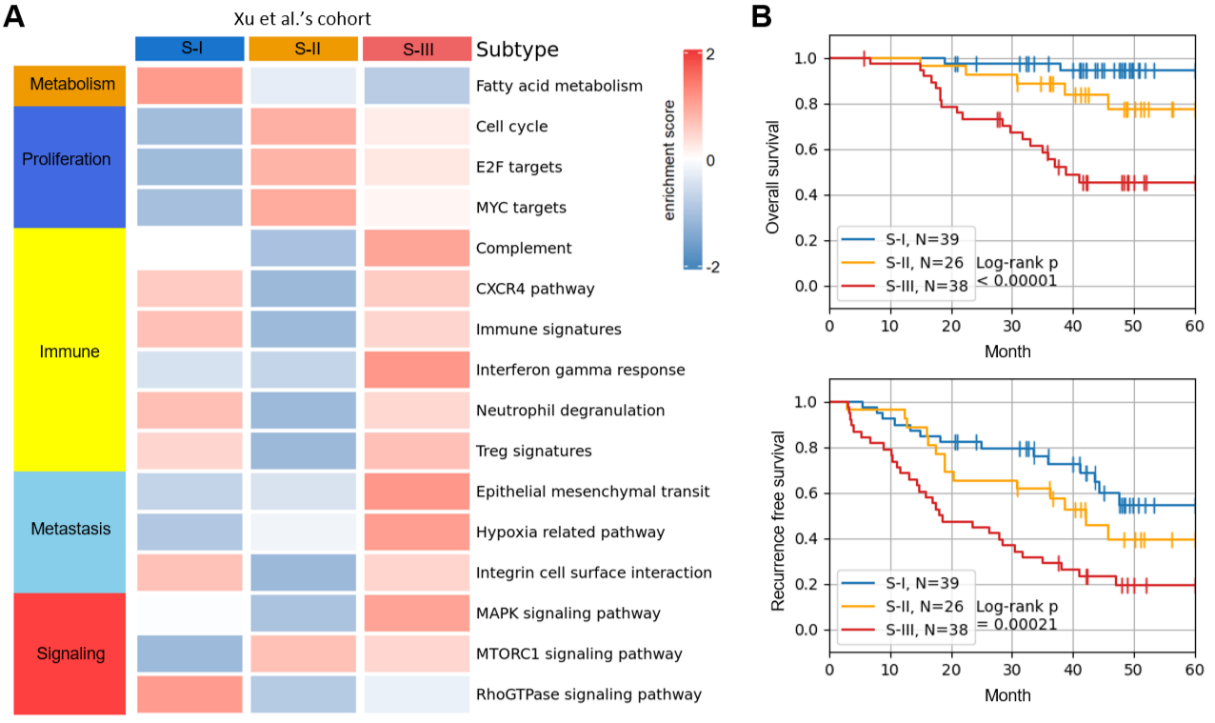

**Figure S8 The visualization of subtyping results on Xu et al.’s cohort**

**A.** The pathway enrichment scores of the SRPS transferred subtypes. **B.** The Kaplan-Meier curves of overall survival and recurrence-free survival of the SRPS transferred subtypes.

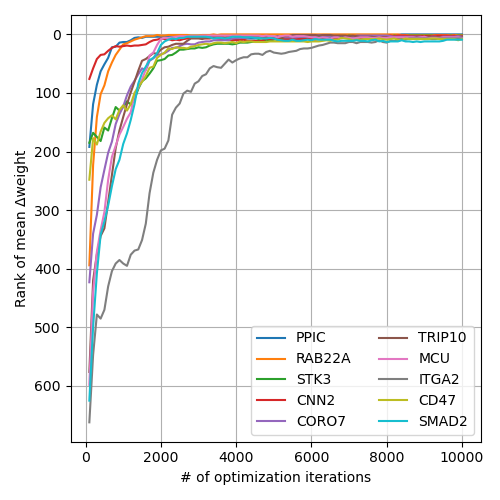

**Figure S9 The rank of protein significance measured by the mean** $\boldsymbol{\Delta}\boldsymbol{weight}$ **converged during the optimization process**

It was calculated based on the averaged $\Delta weight$ of 25 SRPS models (5 folds × 5 repeats). Here, we picked the top-10 proteins in the final models. It can be inferred from the figure that these proteins ranked randomly at the beginning of the training due to the random initialization but finally converged at high ranks. It proves that 10000 steps of optimization are enough to get a stable ranking result.

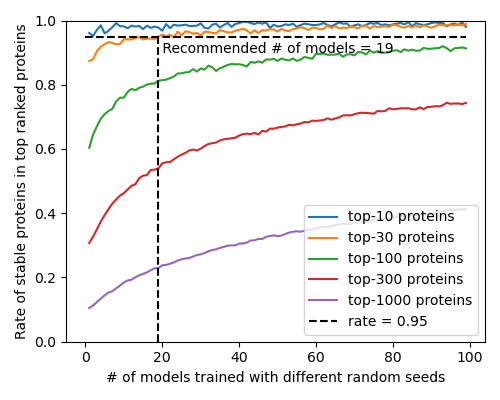

**Figure S10 Stability of the subtype significance score with varying numbers of models**

Curves depict the relationship between the rate of stable rankings in top-K ranked proteins and the number of models used to calculate the subtype significance score. Stability is measured by the consistency of rankings over the last three training epochs across 100 models trained with different random seeds. Each curve corresponds to a specific K value (10, 30, 100, 300, 1000). For top-30 protein analysis, a minimum of 19 models ensures 95% stability in the ranking of subtype significance scores.

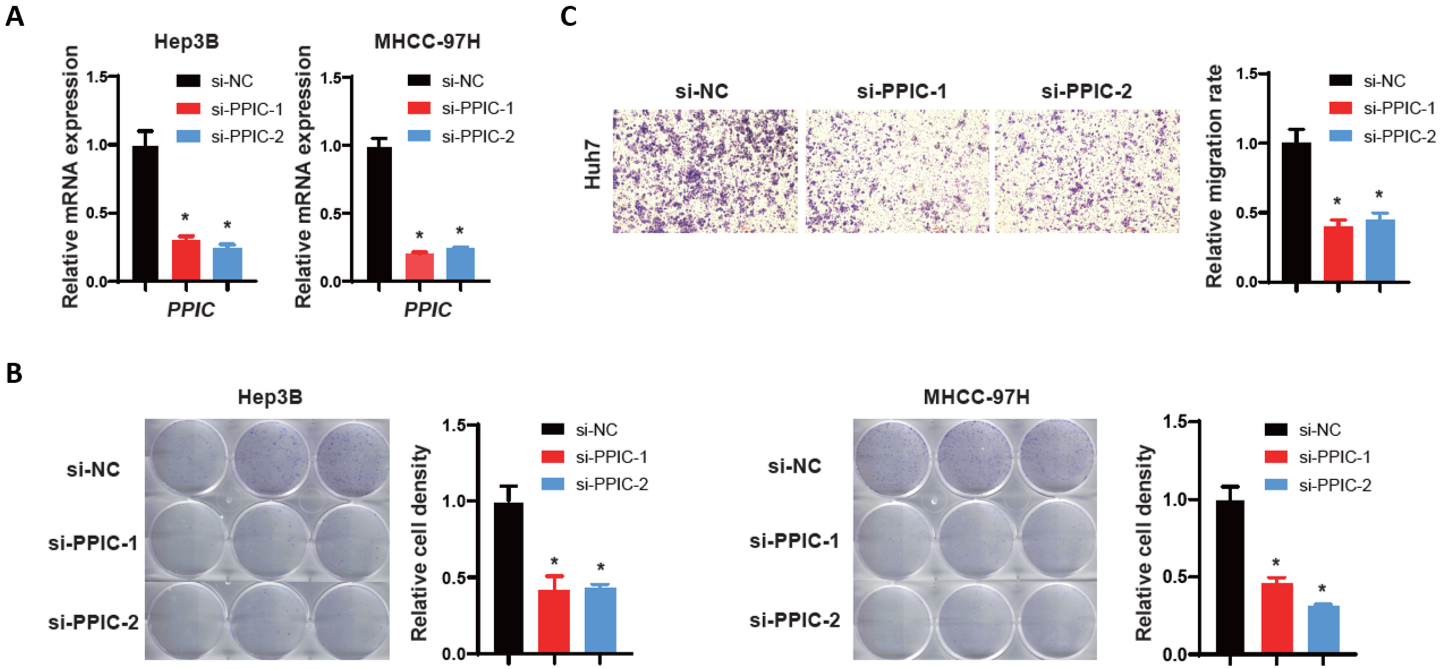

**Figure S11 PPIC promotes cell migration and cell colony formation in HCC cell lines**

**A.** Relative mRNA expressions of *PPIC* after the indicated treatment. **B.** Resulting cell colonies (left) and the related statistical graphs (right) after the indicated treatment. **C.** Resulting staining pictures obtained from cell migration assay after the indicated treatment. Data were presented as mean ± Standard Error of the Mean (SEM). *, ***p***<0.05.

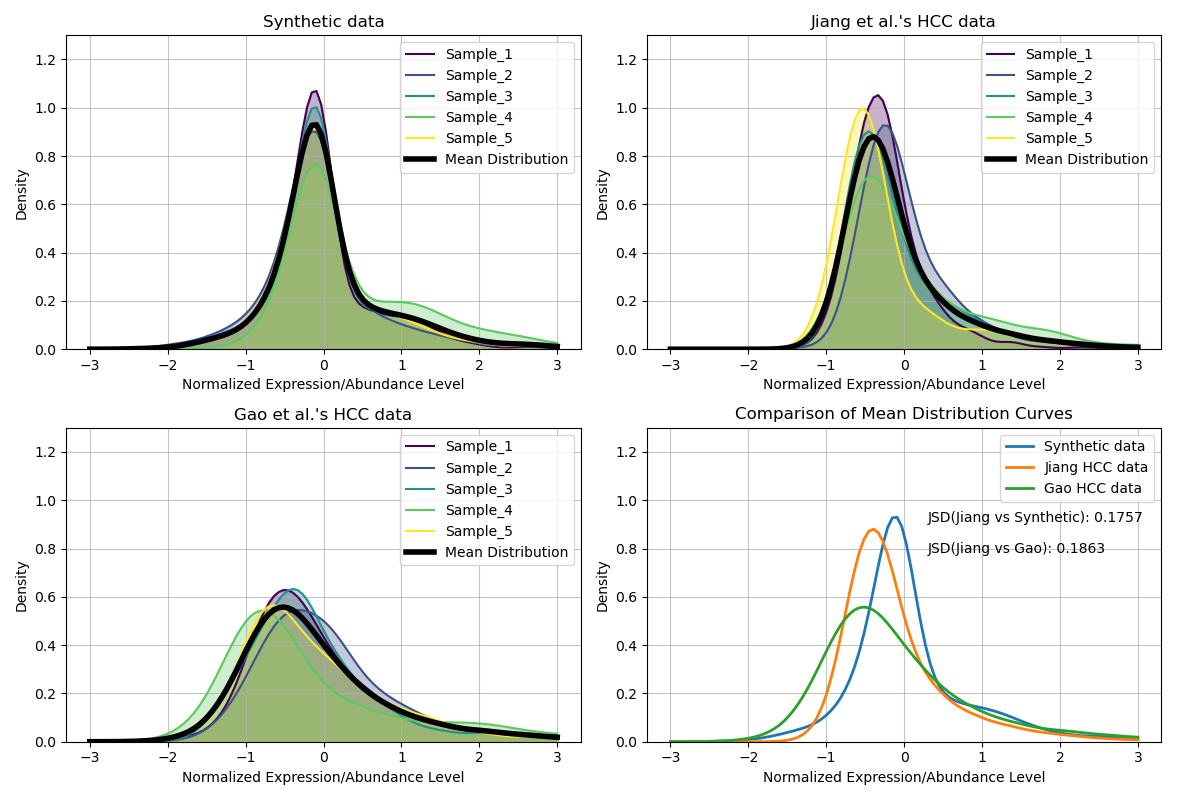

**Figure S12 The distributional divergence between synthetic data and real-world proteomic data**

In this experiment, we randomly sampled data from the simulated dataset (generated using the Splatter package) and two real-world HCC datasets used in our study (Jiang et al.’s cohort and Gao et al.’s cohort). The synthetic data was preprocessed using log2 transformation, and all datasets were further processed with zero-value imputation and Z-score normalization. For each dataset, five samples were selected. The first three subfigures depict the normalized expression/abundance distributions of the samples from each dataset, with the bold black curve representing the average distribution. The final subfigure shows the Jensen–Shannon Divergence (JSD), a symmetric distance metric between distributions, calculated between the average distributions of the datasets. The results demonstrate that the divergence between the synthetic data and Jiang et al.’s cohort is as close as the divergence between the two real-world HCC datasets, which were generated using distinct proteomic quantification methods (Label-Free and TMT-labeling).

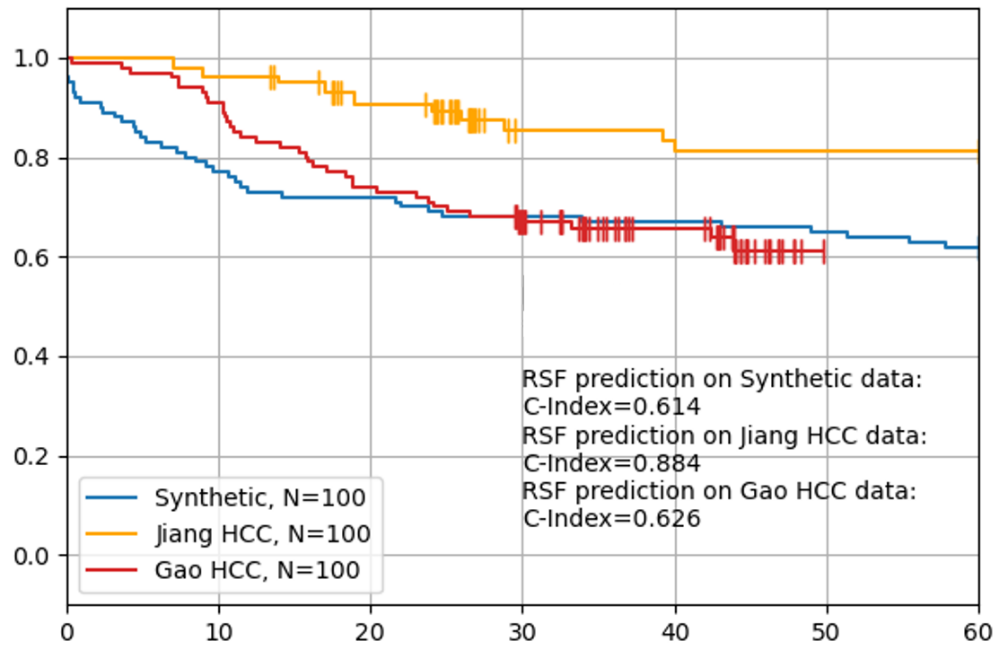

**Figure S13 Comparing the difficulty of the prognosis modeling on synthetic dataset and real-word datasets**

Kaplan-Meier curves show the overall survival distributions for 100 samples randomly drawn from the synthetic dataset, Jiang et al.’s HCC cohort, and Gao et al.’s HCC cohort. The simulated survival curve is comparable to Gao et al.’s cohort. Prognostic performance was assessed by training a Random Survival Forest (RSF) model on each dataset with a 7:3 train-test split. The C-index values are presented, showing similar levels of prediction difficulty between the synthetic dataset and Gao et al.’s cohort.
